# Spatial control of light-responsive proteins and optogenetics within hydrogels via volumetric bioprinting

**DOI:** 10.64898/2026.04.09.717404

**Authors:** Davide Ribezzi, Pere Català, Gabriel Größbacher, Paulina Nuñez Bernal, Olaf Nijssen, Sammy Florczak, Wilco Nijenhuis, Nuria Crusellas-Villorbina, Bram Nijhoff, Paul Delrot, Jos Malda, Andreas Hierholzer, Martin Fussenegger, Lukas C. Kapitein, Riccardo Levato

**Author notes:** Correspondence (R. Levato). These authors contributed equally (Davide Ribezzi, Pere Català).

## Abstract

Spatiotemporal control over cell fate and behaviour within bioprinted constructs remains a key challenge in tissue engineering. Optogenetics offers versatile potential for non-invasive regulation of biological processes. Yet, its integration within large-scale, cell-laden bioprinted materials is still limited, especially considering spatial constraints of existing light delivery methods. In this study, we introduce a novel approach that repurposes tomographic volumetric bioprinting to enable post-printing stimulation of photosensitive protein-switches and optogenetic circuits in cells deep within hydrogel constructs. By converging different bioprinting approaches, computer vision, context-aware model generation, and synthetic biology and cell engineering, we demonstrated selective activation of a fluorescent, light-responsive protein probe within multi-material centimeter-scale constructs. Moreover, leveraging a multi-wavelength volumetric bioprinter, we further demonstrate this concept by selectively stimulating cells expressing a near-infrared optogenetic system that triggers gene expression and the induction of pancreas-specific transcription factors. The described methods provide platforms for remote, repeatable, and localized control of biological events in volumetric constructs, opening new possibilities for advanced tissue models, and dynamic tuning of cell-mediated protein production in engineered living systems.

## 1. INTRODUCTION

Advances in bioprinting technologies have opened new opportunities for tissue engineering and regenerative medicine, enabling the production of complex living constructs that mimic the geometrical and cellular composition of native tissues.^[1,2]^ In particular, recent innovations have significantly advanced the field of biofabrication by improving key aspects, such as resolution and speed,^[3–6]^ expanding design freedom,^[7–9]^ enabling multi-material fabrication and the creation of complex vascular channels,^[10,11]^ and enhancing control over porosity,^[12–14]^ and biochemical gradients.^[15,16]^ Yet, for bioengineered tissues to acquire *in vivo*-like functions, bioprinting is only the first of many steps, and must be followed by post-printing maturation.^[17]^ To date, post-printing control of cellular events is typically induced with molecules added to the media used for tissue culture. This approach offers poor control over the spatio-temporal delivery of the stimuli and is detrimental when effects are desired only in one cell type or region. Including controlled delivery devices (i.e., micro- and nanoparticles) to release bioactive molecules within a printable material can mitigate the problem,^[18,19]^ though it does not overcome off-target effects due to diffusion. Elegant strategies were also proposed using drug-triggered gene expression in engineered cells to induce cell differentiation,^[20]^ though the stimulus cannot be turned off on demand.

Considering these challenges, light stimuli are especially beneficial, as they influence cells in a non-invasive and biocompatible manner, upon optimization of the light dose, and reactive species concentration.^[21–23]^ As such, light-responsive chemistries have been often investigated to facilitate control of cell-materials interactions, and presentation of bioactive stimuli in tissue engineering.^[24,25]^ A rising number of light-responsive proteins and molecules have been discovered, each reactive to different UV, visible, and infrared wavelengths, often orthogonal to the light commonly used for printing hydrogels.^[26–28]^ Moreover, light-responsive proteins have also been used to build smart hydrogels with shape-changing capacity,^[29]^ enzymatic functions,^[30]^ and to activate materials-embedded dormant growth factors,^[31]^ therefore, potentially offering a broad array of tools to interact with cells in biofabricated constructs. Notably, many light-controllable molecules are part of the toolkit of optogenetics, which encompasses techniques for fast and precise control of biological systems upon illumination.^[32–35]^ In addition to applications for fundamental research, designer cells equipped with optogenetic circuits have demonstrated therapeutic effects in animals, for example by restoring reproductive health,^[33]^ by managing glucose homeostasis in experimental diabetes models,^[36]^ and even in humans, in the context of vision restoration.^[37]^ Recently, our team has also shown that optogenetic circuits integrated into bioprinted constructs can be used to regulate cell behavior, as shown via blue light-mediated insulin release from engineered pancreatic cells.^[7]^ Nevertheless, to date, spatial-selective photochemical control of light-sensitive proteins and optogenetics experiments have been either performed using 2D projections (i.e. created by (photo)masks,^[34,38]^ or focused beam as is the case of light-sheets^[39]^) which provide no control in the direction of propagation of the light, as every cell along the light path is activated, or in 3D, using two-photon illumination systems.^[40,41]^ Multiphoton effects, however, although providing subcellular resolution, are limited to a working distance typically shorter than a few hundreds of µm.^[41]^ This hampers the targeted 3D stimulation of voxels of interest within large objects and, therefore, limits the ability to effectively be used in physiologically-relevant sized tissue and biomaterial-based constructs. Notably, pinpointing or selecting areas deep within cell-laden structures of interest remains not possible. This effectively precludes the possibility to build compartments, gradients, and anisotropic patterns or optogenetic activation, in which the stimulated region is truly confined in x-y-z across tissue constructs of sizes relevant for many applications in tissue engineering and, organ-on-chip, and advanced *in vitro* human disease models.

Addressing this key gap, in the present study, we introduce a new solution to regulate cellular events across desired regions within-centimeter scale hydrogel constructs. We propose that light inducible proteins and optogenetic circuits can be controlled with spatial precision at any time point post-printing, leveraging advanced light-based bioprinting devices via tomographic volumetric bioprinting. Insofar, volumetric printing has been used to perform photochemical reactions and building large-scale objects in a layer-less fashion, utilizing single-photon, visible light tomographic projections.^[42–44]^ Herein, by extending the use of tomographic light doses on cellular processes, the same light input can be used to control the biological events or function of the construct at any time point with precise spatial pinpointing and 3D control. Leveraging the capacity of tomographic printing to operate with a broad range of wavelength and with multi-colour light systems,^[8]^ we demonstrate the activation of diverse photosensitive proteins. As the first proof-of-concept, we first focused on cells expressing a reporter light-sensitive protein, which undergoes a fluorescence bandwidth shift once exposed to blue light. We produced several volumetrically printed designs and showed how targeted tomographic delivery correctly induces region-selective activation of the reporter. Of note, the reactivity of photosensitive proteins can typically be modulated by the delivered light dose, offering an additional layer of control and the possibility to introduce gradients of degrees of activation. Moreover, combining the idea of tomographic activation of photosensitive proteins with the recently described concept of context-aware 3D printing,^[11]^ we use printer-paired imaging and computer vision workflows to map and detect specific cell clusters, and provide targeted protein photoswitch of selected cell-laden regions. Finally, we provide a proof-of-principle experimental evidence of the possibility to stimulate optogenetic gene expression circuits, using cells expressing a near-infrared responsive photoreceptor. This technology opens new avenues for controlling cell-laden hydrogels, bioprinted tissues and engineered living materials, offering a powerful tool for developing *in vitro* models for biomedical research.

## 2. RESULTS AND DISCUSSION

### 2.1 Generation of cells carrying a photoswitch protein probe

To first assess the possibility to perform tomographic activation of photosensitive proteins, we generated a stable HeLa cell line expressing the photoswitchable protein mEos3.2. mEos3.2 is a naturally green fluorescent (λ_ex_ 505 nm) protein which undergoes photoconversion when irradiated with 405 nm light, leading to a widening and shifting of the absorption spectrum, resulting in the acquisition of fluorescence in the orange region of the visible spectrum (λ_ex_ 590 nm).^[45]^ When photostimulated, cells carrying mEos3.2 display fluorescence both in the green and orange region of the visible spectrum. The use of such a protein offers a rapid and easy readout through light intensity and channel selection (**Figure 1A**). Expression of the protein is linked to a tetracycline operator, to avoid unwanted expression and photoconversion during the cell expansion phase ahead of bioprinting and of optogenetic stimulation experiments. After the successful generation the stable HeLa mEos3.2 cell line, we first assessed the feasibility of the photoconversion in a 2D setting, using a digital micromirror device (DMD) mounted to control light patterning onto a microscope. These experiments showed efficient and precise (down to single cell) photoconversion from 507/515 (λ_ex_/ λ_em_) to 572/580 (λ_ex_/ λ_em_), therefore, confirming the *in situ* functionality of the engineered protein construct (**Figure 1B**).

**Figure 1.**
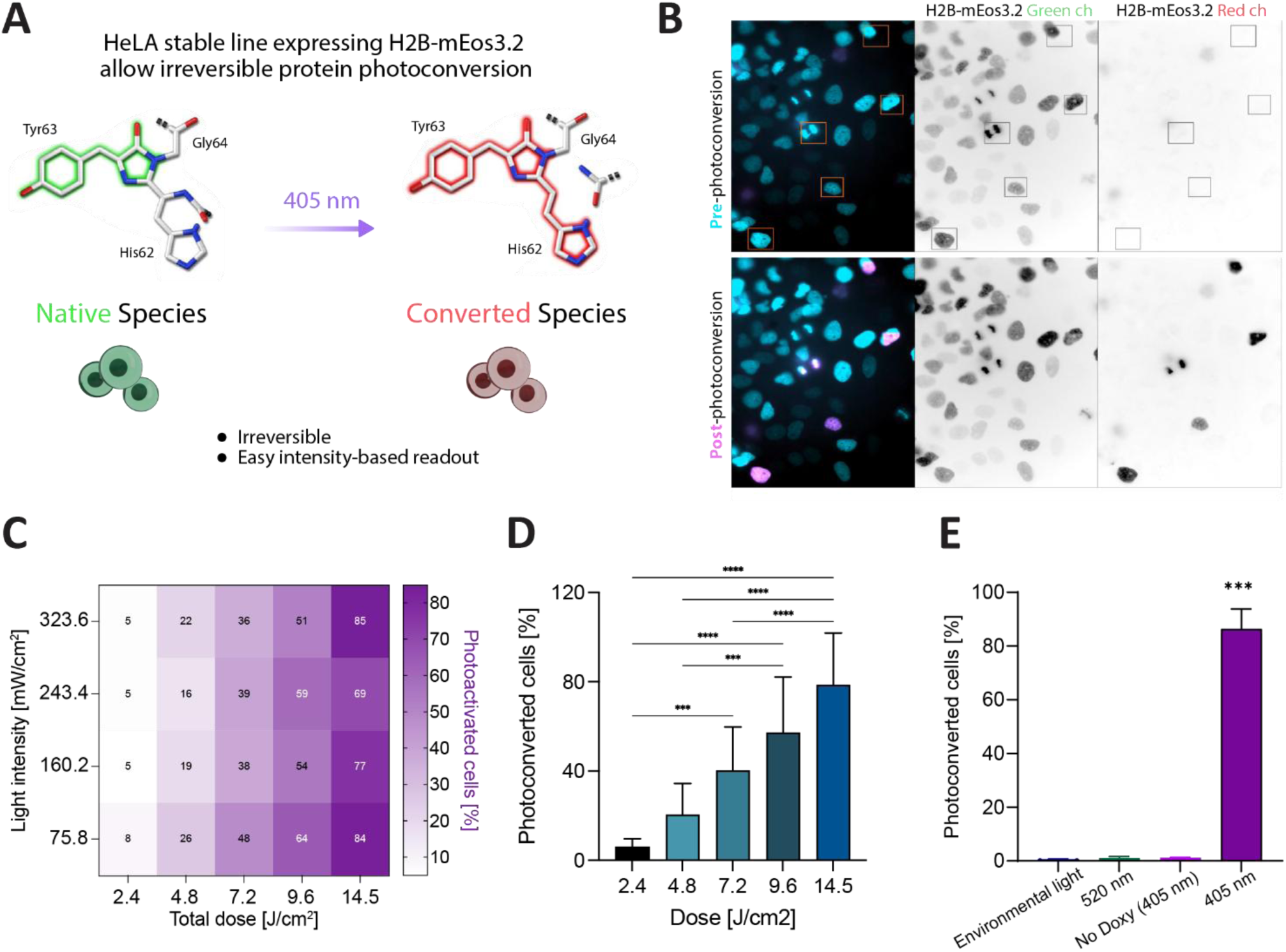
Light-triggered photoconversion and dose-response characterization of the photoconvertible protein mEos3.2 stably expressed in HeLa cells. A) Schematic representation of mESO3.2 conformational change upon 405 nm light irradiation. Adapted with permission from the Journal of Cell Science.^[51]^ B) Photoconversion of mEos3.2 stably expressed in HeLa cells by using a cell targeting photomask (yellow squares represent irradiated cells with 405 nm light). Left: cyan is 470 nm excitation signal and magenta is 590 nm excitation signal. Middle image: 470 nm excitation signal. Right image: 590 nm excitation signal. Images taken at 40x magnification. C) Heat map representing percentage of photoactivated cells after exposure to increasing doses (2.4-14.5 J/cm^2^) and increasing light intensities (75.8-323.6 mW/cm^2^) using a 405 nm laser (n=3). D) Percentage of photoconverted mEos3.2-expressing cells by increasing light-dose exposure (n=12). E) Wavelength specificity of mEos3.2 photoconversion (n = 4). Data represent mean ± SD. Significance was determined by one-way ANOVA (D) or unpaired t-test (E) *** = p <0.001, **** = p< 0.0001.

A key principle in volumetric printing is that photochemical, and by extension photobiological effects, need to display a thresholding behavior, in order to be selectively triggered with single photon tomographic light projection.^[5,46,47]^ Thus, we exposed cells carrying mEos3.2 to different 405 nm light doses, and characterized such thresholding behavior (**Figure 1C** and **Supplementary Figure S1A-B**).

Results showed an increased photoconversion percentage of mEos3.2 proportional to the light dose delivered, irrespective of the specific combination of exposure time and light intensity selected (**Supplementary figure S1B**), with minimal activation at the lowest dose (<6% at 2.4 J/cm^2^), and reaching ⁓80% of photoconverted cells with a 14.5 J/cm^2^ delivered dose (**Figure 1D**). Dose-dependency is also in line with the previously reported data for mEos3.2 kinetics of conversion.^[45]^ Moreover, mEos3.2 showed wavelength selectivity and doxycycline-related stability, resulting in <1% of photoconverted cells upon environmental light exposure and a 14.5 J/cm^2^ light dose delivered with a non-activating wavelength (520 nm); or illuminating HeLa cells cultured in the absence of doxycycline with a 405 nm light at the highest dose value (**Figure 1E**).

Considering the gathered data, HeLa mEos3.2 resulted in a promising platform to be used for testing the hypothesis that bioprinting could be re-routed to provide spatial selective, light-based triggering of biological reactions. Towards this goal, it is important to underline that light at 405 nm is also typically used to trigger free-radical polymerization of bioresins often used in light-based bioprinting techniques, mostly due to the widespread use of lithium phenyl-2,4,6-trimethylbenzoyl-phosphinate (LAP) as a photoinitiator.^[48,49]^ Nevertheless, our dose-response analysis (**Supplementary Figure S2**) identified that the minimum light intensity needed to obtain a photo-conversion of mEos3.2 exceeded 10 times the 0.18 J/cm^2^ dose used for cell encapsulation, a value in the range of typical light doses reported in volumetric bioprinting at 405 nm,^[5,50]^ suggesting it is possible to sequentially perform both photochemical processes in an independent manner.

### 2.2 Sequential volumetric bioprinting and spatial selective mEos3.2 photoconversion within complex 3D constructs

After generating the stable HeLa mEos3.2 cell line and having characterized its responsiveness to light stimuli in 2D, we assessed the feasibility to trigger the biological photoconversion using the laser of the tomographic volumetric printer, and analysed the dose-response behaviour of the cells also when embedded in a hydrogel bioresin, at first, using 5% w/v GelMA, supplemented with 0.1% w/v LAP as photoinitiator, as platform material for testing. Also in 3D, the dose needed to activate the photo-switching protein is higher than that used for photocrosslinking (Supplementary Figure S2), which allowed us to perform the alignment procedure with the green laser and two reactions (printing and photostimulation) independently and sequentially. We observed the presence of a threshold effect for mEos3.2 photoconversion also when using the volumetric printer on hydrogel-embedded cells. A light dose >5.0 J/cm^2^ was required to observe photoconverted cells, with activation efficiency increasing at higher doses, independently of the chosen light intensity, underlining how the conversion is primarily dose-dependent also in 3D samples (**Figure 2A**). In particular, a spatial activation fidelity (the ability to accurately replicate the photomask pattern onto the hydrogel with HeLa activation) of 90 ± 8%, with an efficiency of 84 ± 7% of cells converted, was reached with the GelMA-based bioresin. In parallel, we also confirmed the photoconversion process in 3D could be applied also to different types of hydrogels, using Alginate Methacryloyl (AlgMA) as a polysaccharide-based bioresin (**Supplementary Results, Supplementary Figures S3-5**).

**Figure 2.**
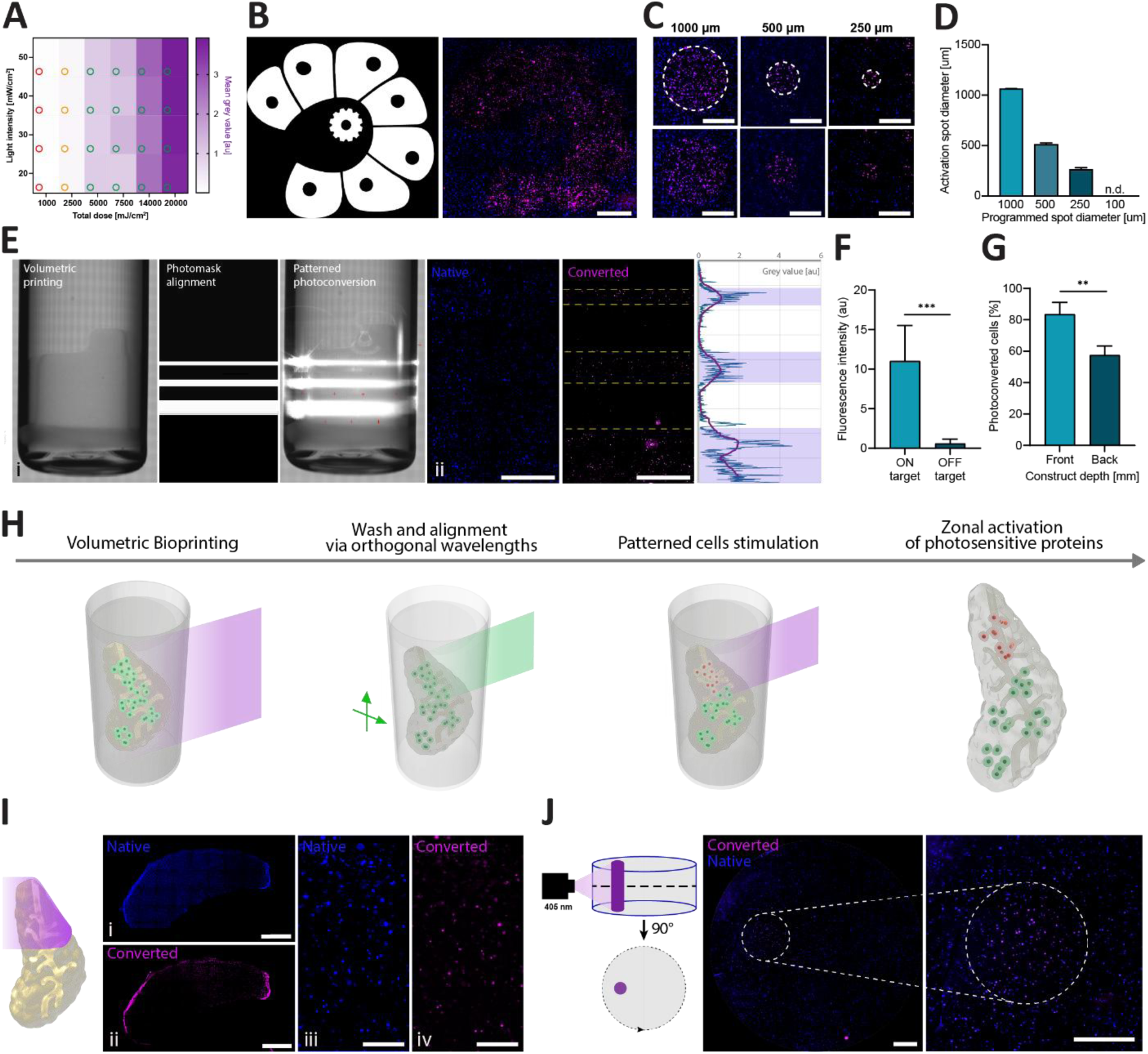
Spatial selective mEos3.2-expressing HeLa cell photoconversion in volumetric bioprinted constructs. (N = native HeLa-mEos3.2; P = photoconverted HeLa-mEos3.2). A) Screening of volumetric printing set-up light intensity and total light dose required for HeLa-mEos3.2 photoconversion. Red circles represent exposed regions that did not result in mEos3.2 photoconversion and green circles represent regions where photoconverted HeLa-mEos3.2 were present. B) Activated HeLa-mEos3.2 cells using a photomask of our laboratory logo. Measurements of the size of the smallest (negative) features are reported in the main body of the manuscript (n =3). The left panel shows the projected photomask, the right panel shows the fluorescent microscopy image. Magenta represents activated/photoconverted HeLa cells, while blue represents non-converted HeLa cells. Scale bar: 1 mm. C) Resolution screening of photoactivated regions using an array of dots with varying size. Close-up images of activated circular patterns of 1000, 500 and 250 μm diameters after exposure to 20000 mJ/cm^2^,are showed and D) the diameter of the photoactivated regions is quantified (n = 3). Scale bars: 500 μm. E) Camera view of (i) the volumetric printing vial inside the bioprinter containing the prism (left, volume = 140 mm^3^), photomask alignment (middle), and photostimulation process (right) (ii) Fluorescence quantification of the non-photoconverted HeLa cells (right) and the photoconverted HeLa cells (middle), including a plot fluorescence profile of the targeted and non-targeted regions (left) (magenta line representing the median of the blue plot). Scale bars: 1mm. F) Greyscale values of the targeted and non-targeted regions, as observed in the 590 nm channel post photoactivation. G) Visualization of the photoconversion efficiency attenuation throughout the construct thickness, when the rotating platform holding the sample vial is maintained still (n = 4). H) Schematic representation of the workflow of the sequential volumetric bioprinting and photo-conversion process. I) 3D photoconversion of HeLa mEos3.2 in a bioprinted pancreas-shaped model (volume = 250 mm^3^) including a i,ii) fluorescence tilescan of the whole construct. Scale bars: 2 mm. iii,iv) Core of the construct. Scale bars: 250 µm. J) Tomographic photoconversion of an offset 1-mm cylinder within a printed GelMA cylinder (volume = 193 mm^3^). Scale bars = 500 µm. Data represent mean ± SD. Significance was determined by unpaired t-test (F, G): ** = p <0.01, *** = p <0.001.

Notably, using the DMD of the printer also permits to deliver more complex light patterns directly onto bioprinted constructs. To showcase this, we initially projected the image of our lab logo of the printer to produce a static photomask (**Figure 2B**), effectively triggering mEos3.2 photoconversion in a highly spatially-selective manner, as the activation pattern follows the intended design of the projected logo, in which the smallest features, the holes in the petals surrounding the cog measure 397.8 ± 29.9 µm (intended diameter = 402.5 µm, and the hole in the center of the cog measures 283.3 ± 11.3 µm (intended diameter = 292.9 µm).

Next, we further characterized the limits of resolution that could be achieved with the current hardware of the printer, by delivering circular patterns of decreasing size, and assessing the size of the region with activated cells. Even when using the highest value indicated in the dose test (20 J/cm^2^), which could easily activate cells off-target if not delivered correctly, we found good pattern fidelity (with 1064.9 ± 4.3, 512.3 ± 15.2, 262.7 ± 19.2 µm for the 1000, 500, and 250µm circles, **Figure 2C-D**). Notably, the smallest circle in which activated cells could be reliably found measured 250µm in diameter, while when setting the delivered circle to 100 µm, no identifiable spot with photoactivated cells could be recognized. In addition, it should be noted that resolution of the volumetric photomanipulation could be further enhanced via adopting hardware solutions for volumetric tomographic printing that enable higher spatial resolution, as recently shown with the printing of features in the range of 10µm, using holographic light fields.^[52]^

Subsequently, we explored the possibility to trigger the photoconversion in larger-scale 3D hydrogels. As a first demonstration, we bioprinted and anchored to the vial a prism (5 x 5 x 5 mm, with a taller side to distinguish the front from the back of the construct for subsequent analysis) embedding HeLa mEos3.2 at 10^6^ cells/mL concentration, which was stimulated with light projected following a with reference features (three parallel lines with different width, 250, 500 and 1000 µm) (**Figure 2Ei**). Photoconverted HeLa mEos3.2 were found only within the targeted regions (**Figure 2Eii**) (with an activation accuracy of 84 ± 15%, 92 ± 7%, and 96 ± 3% respectively for the 250, 500 and 1000 µm bands) and significantly higher fluorescent intensity values could be measured from the targeted regions (11 ± 4 au) compared to the non-targeted ones (1.6 ± 1.3 au) (**Figure 2F**).

In 3D samples, the light absorbance from the material and the scattering caused by the cells within the volume can alter the optical properties of hydrogel (**Supplementary Figure S6**), challenging the light penetration and focus within the volume.^[53,54]^ After measuring the photoconversion efficiency at the back of the prism, we indeed noticed a significantly lower percentage of photoconverted cells (57 ± 4%) compared to the front (83 ± 7%) of the construct (**Figure 2H**). It should also be noted that, while these preliminary experiments were performed with static light projections onto the hydrogels, tomographic volumetric bioprinting, being performed on a full rotation around the target material, could also mitigate these differences. Therefore, as a proof-of-concept of the sequential volumetric bioprinting and 3D photoconversion process for the generation of engineered constructs, we first encapsulated the HeLa mEos3.2 cells in a printed pancreas-shaped mode, in which only the top half was subsequently photostimulated (**Figure 2H**). This led to the successful, spatially confined photoconversion of the mEos3.2 only in the illuminated region (**Figure 2Ii**) with no detectable variation in the number of converted cells across the stimulated volume (**Figure 2Iii**). In addition to that, a lymph node-inspired geometry, containing perfusable channels simulating the lymphatic vessels perfusing the native tissue was volumetrically bioprinted, in which specific regions simulating the native germinal centres were selectively photoconverted (**Supplementary Figure S7**). Besides the possibility to combine printing with optogenetic stimulation, a unique feature of volumetric printing is the capacity of using single-photon light sources to pinpoint anisotropic light doses inside hydrogels and not just along the projected light path. Showcasing this ability, tomographic reconstructions were used to successfully resolve a 1 mm diameter cylinder of activated cells inside a cell-laden hydrogel disk (**Figure 2J**), a feature that opens the possibility to confine optogenetic proteins activation selectively within an engineered tissue, while limiting or preventing unwanted activation along the path of the incident light beams.

### 2.3 Embedded Extrusion Volumetric Printing for spatial-selective photoconversion within multi-material 3D constructs

Having showed how VBP can be used for photostimulatation in constructs comprising cells homogenously suspended throughout the volume, next, we investigated how to achieve spatial-selective photoconversion within high-cell density features patterned inside hydrogels. For this, we used an approach based on the recently described Embedded Extrusion Volumetric Printing (EmVP) process,^[7]^ which is especially beneficial when aiming to position multiple cell types or materials in a volumetric bioprinting set-up (**Figure 3A**). The embedded extrusion step of HeLa mEos3.2 cells was performed in a low molecular weight GelMA-based fluid bulk support bath, optimal for its self-healing-like properties, as previously reported,^[8]^ followed by volumetric printing to sculpt the construct in its final, crosslinked shape, and eventually via subsequent targeted activation of the photoswitching protein. A characterization of the rheological properties of the fluid bath is reported in **Figure 3B-D**. Following the printability optimization (**Figure 3E**), filaments of width ranging from 331 ± 23 µm to 134 ± 22 µm could be obtained via extrusion of cells carried by a sacrificial, water soluble methylcellulose-based ink, which was previously designed and characterized^[6]^. As a proof-of-concept of the integration of EmVP with specific spatial photoconversion of the mEos3.2 construct, we generated a structure consisting in three extruded rings of clustered HeLa mEos3.2 cells, embedded into a low MW GelMA based cylinder scaffold with a central channel (**Figure 3F**).

**Figure 3.**
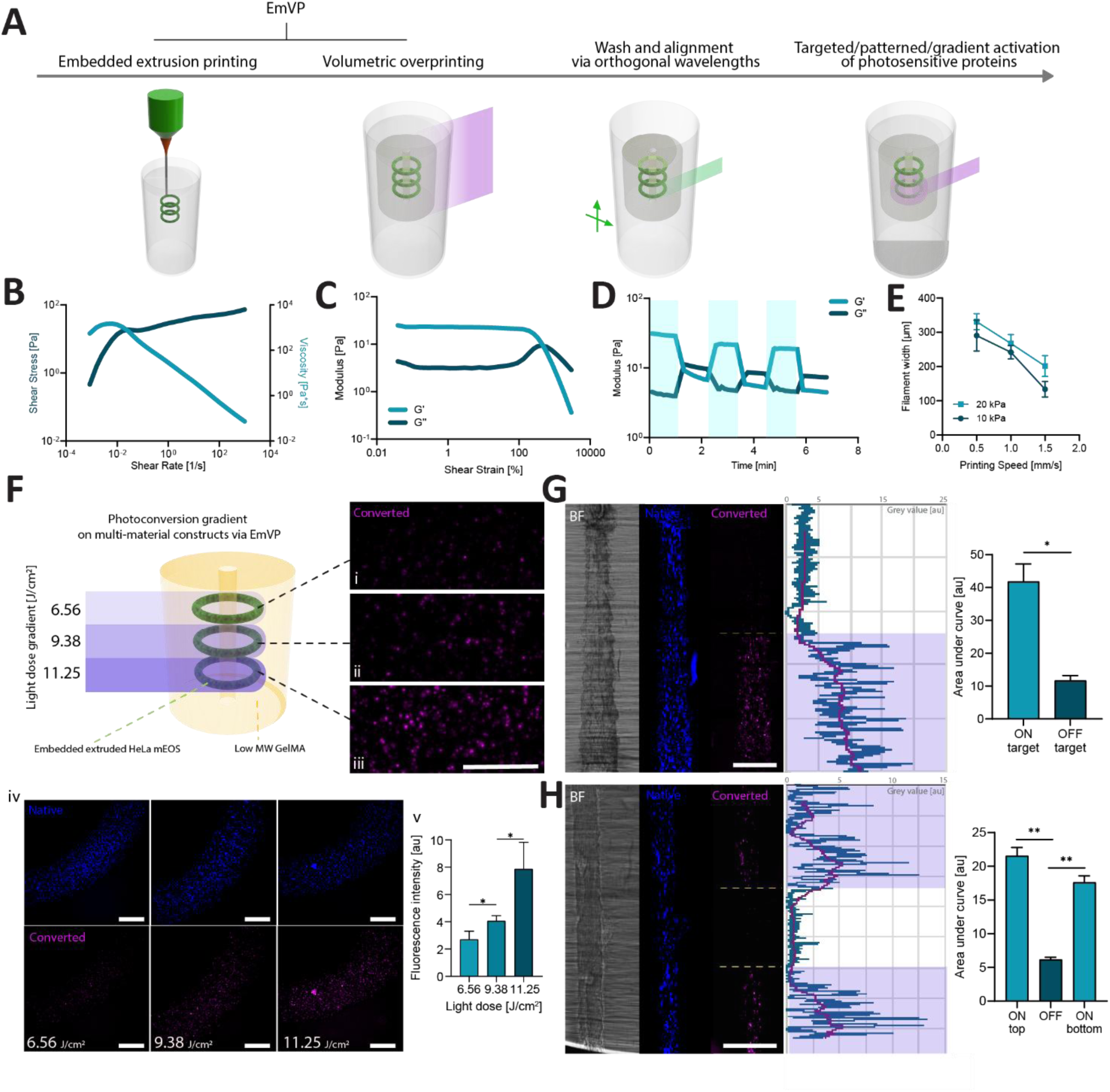
Photoconversion gradients on complex multi-material constructs via EmVP. A) Workflow of the EmVP and HeLa-mEos3.2 photo-stimulation process. B-D) GelMA fluid bulk support bath representative graphs (n = 3) of the rheological properties that make it suitable as suspension bath for embedded extrusion bioprinting. B) Shear thinning behaviour observed as the viscosity decreased as the shear strain increased and, under the same conditions, the shear stress increased in a nonlinear fashion. C) Shear-yielding with increase in strain (0.037–1000%, 1 Hz). D) Stress/recovery through low (unshaded, 1% strain, 1 Hz) and high (shaded, 500% strain, 1 Hz) strain cycles. E) Printing accuracy of the 2% w/v methylcellulose bioink, calculated by measuring the filament width as a function of the extrusion pressure and printhead translational velocity (n = 6). F) Photoconversion gradients on complex multi-material constructs produced via EmVP (n = 3) of a total volume of 190 mm^3^. Scale bar is 500 µm (iv) G-H) Bright field and fluorescent images of the native and converted cells in extruded filaments (produced within a construct with overall volume of 220 mm^3^) stimulated with light patterns. Scale bar is 1 mm. Values for the area under curves (normalised by the illuminated length of the filament) were obtained from the intensity profile plot of the targeted regions, compared to the non-targeted ones (n = 2). Data represent mean ± SD. Significance was determined by unpaired t-test (F, G, H) * = p<0.05, ** = p<0.01.

Using the DMD of the printer, we generated a light intensity gradient at 405 nm targeting the three extruded rings, resulting in three different delivered light doses for each ring (6.56, 9.38 and 11.25 J/cm^2^ dose intensities for the top middle and bottom ring. **Figure 3Fi, ii** and **iii**, respectively). Results showed the possibility to create activation gradients on multi-material constructs, where highly dense extruded features (containing 30 × 10^6^ cells/mL) targeted with a higher stimulating dose resulted into a significantly higher fluorescence signal, leading to fluorescent intensity values of 2.7 ± 0.6 au, 4 ± 0.3 au and 7.8 ± 2 au for the top, middle and bottom ring, respectively (**Figure 3Fiv** and **v**). While the light doses we used appeared to be non-harmful for HeLa cells, it could not be excluded that the highest dose could lead to phototoxicity to more sensitive cell types, such as primary, tissue-derived cells. Therefore, we also assessed viability of encapsulated cells upon additional illumination to doses up to above the highest tested in these experiments, using bone marrow derived mesenchymal stromal cells of equine origin, confirming no negative effect on viability (**Supplementary Figure S8**). Nevertheless, as different cell types could be differentially sense light-mediated stresses, it is recommended to screen compatibility of the photoillumination protocols every time using a new cell source is needed.

Additionally, we tested the possibility to shine light patterns on a single extruded component to recreate accurate on/off activation zones. To do this, we extruded HeLa mEos3.2 cells into straight filaments, embedded them into 3D structure with EmVP, and subsequently delivered 405 nm (11.25 J/cm^2^) light bands to the filaments. Results showed it was possible to photostimulate half of a 5 mm-long filament (**Figure 3G**). This observation was quantitatively confirmed via image analysis, drawing an intensity plot profile along the filament. Measuring the area under the profile curve showed a value of 41.8 ± 5 au for the exposed area compared to 11.7 ± 1 au for the unexposed area.

Activation patterns of bands with a 1.5 mm gap also resulted in precise and high-contrast activated and unactivated regions, measuring an area under the curve of the plotted intensity profile of 21.5 ± 1 au and 17.6 ± 1 for the top and bottom illuminated band, respectively, compared to a significantly lower area of 6.4 ± 0.3 for the non-exposed region. Additionally, we tested the possibility to shine light patterns on a single extruded component to recreate accurate on/off activation zones. Cell filaments embedded into 3D structures via EmVP were activated in a spatially selective manner, first by photostimulation of only half length of a 5-mm filament (**Figure 3G**). This observation was quantitatively confirmed via image analysis, drawing an intensity plot profile along the filament, as measuring the area under the profile curve showed a 3.7-fold significant increment in fluorescence intensity over the inactivated baseline. Activation patterns of bands with a 1.5 mm gap also resulted in high-contrast activated and non-activated regions (**Figure 3H**), measuring an area under the curve of the plotted intensity profile of 3.4-fold and a 2.8-fold significant increment over non-exposed regions for the top and bottom illuminated band, respectively. As indicator of resolution and spatial fidelity, we also observed that length of the activated portions of the filament was 7.0 ± 2.3 % longer than the intended design, likely due to cell-mediated scattering, suggesting that for even more precise photostimulation, this minor deviation should be taken into account when designing the projection masks.

### 2.4 Highly selective and feature-driven volumetric bioprinting and 3D photostimulation

For light-driven 3D-localized stimulation of precise cell clusters, strategies for the identification and subsequent automated tracking of cells or regions of interest that need to be targeted within a tissue are required. Recently, we introduced a novel approach termed generative, adaptive, context-aware 3D printing (GRACE) that uses computer vision and parametric design criteria to enable volumetric printers to create and produce print designs, that conform to the position of cells within a construct, such as vascular channels designed to reach every cell cluster in a volume (**Supplementary Figure S9-10**).^[10]^ This technique could be particularly helpful in identifying cells to be illuminated, and perform context-aware driven photoconversion within a bioprinted structure. Therefore, we designed an additional step to the GRACE workflow, aiming at a highly-specific photoconversion of the encapsulated cells, even in an obstacle-crowded environment. Firstly, we set up a coaxial nozzle-based system for the generation of alginate microspheres encapsulating HeLa mEos3.2 cells at a high density (**Figure 4A**). A 2% w/v alginate solution was selected as bioink for microspheres generation, being this a well-established and characterized biomaterial in the biofabrication field, and offering a fast and simple physical crosslinking process, optimal for the generation of microspheres and cell encapsulation.^[55–57]^ With a set extrusion pressure level between 20 - 30 kPa (minimum needed to extrude the bioink) and varying the air pressure to break the extruded filament and generate particles, results showed how microsphere diameters decreased when increasing the pressure in the coaxial air flow, down to 121.6 ± 40 µm for 120 kPa (**Figure 4B**). To reach a high number of cells within a single particle, we used pressure values at the lower end of the investigated range (20 kPa and 30 kPa for bioink extrusion and airflow, respectively) to generate mm-scale spheres to test in the GRACE workflow. The encapsulation process was safe for the cells, resulting in high viability (89.2 ± 0.5%) after extrusion (**Figure 4C**) and up to 7 days post-encapsulation (**Supplementary Figure S11**). Next, we assessed the feasibility to photoswitch HeLa mEos3.2 post encapsulation. By delivering the same light dose values of 11.25 J/cm^2^ used to stimulate the features in the EmVP process, we observed a successful photoconversion (**Figure 4D**), resulting in 85.3 ± 2% of activated cells (**Figure 4E**).

**Figure 4.**
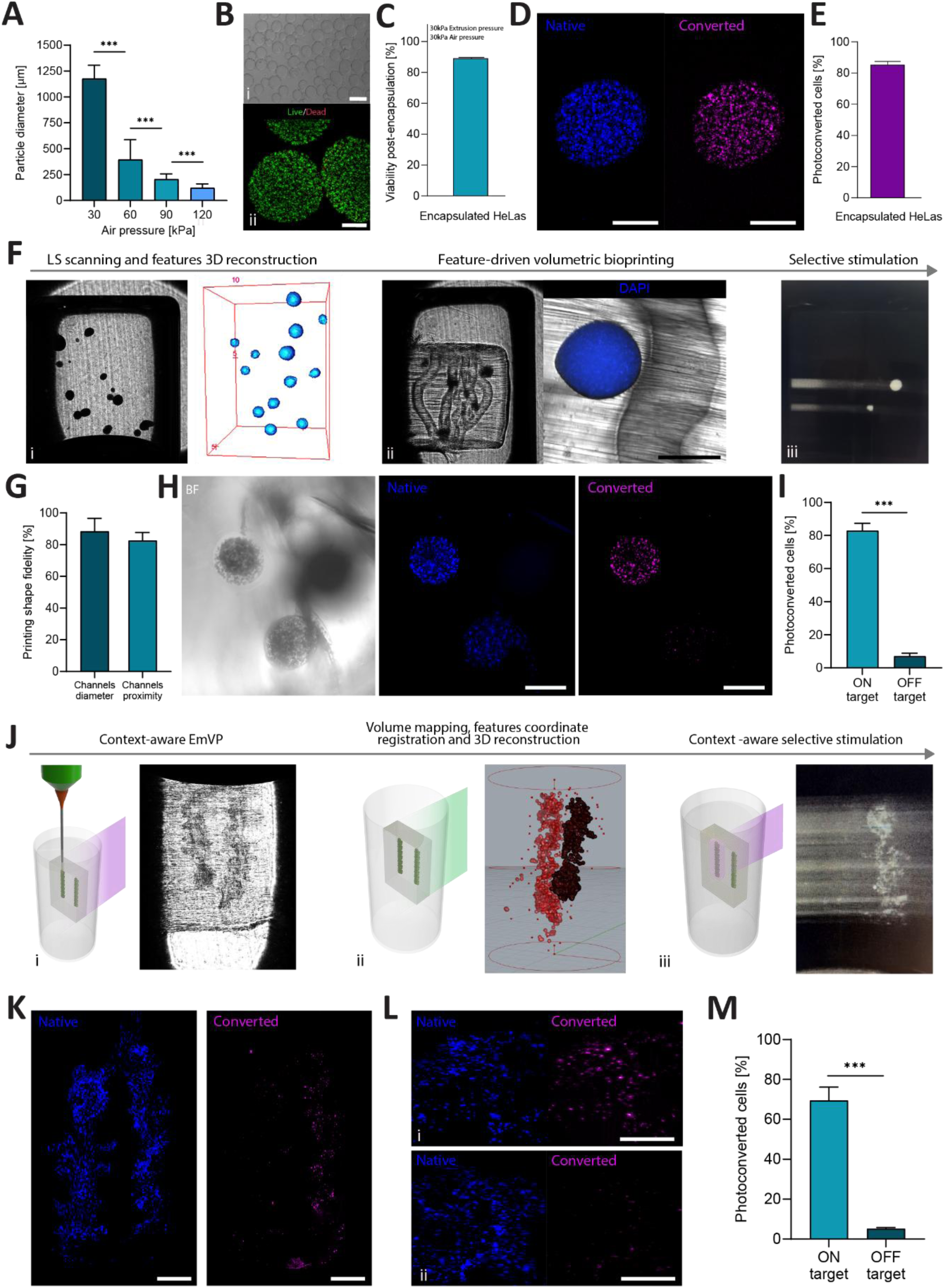
GRACE allows printing adaptive and feature-driven models with complex geometries and selective photostimulation of encapsulated HeLa mEos3.2 cells. A) Tuning of alginate microspheres size by adjusting air pressure during air-assisted coaxial extrusion. Hydrogel extrusion pressure was kept between 20 - 30 kPa, while air pressure varied from 30 kPa to 120 kPa (n = 60). B) (i) High throughput production of low polydispersity alginate microspheres. Scale bar: 200 µm. (ii) Live/Dead staining of HeLa mEos3.2 immediately after the encapsulation process. Scale bar: 500 µm. C) Viability % of HeLa mEos3.2 cells after encapsulation with 30 kPa for both air and extrusion pressure (n = 4). D) Close up of the alginate microsphere embedding HeLa mEos3.2 cells in their native and converted state. Scale bar: 500 µm. E) Encapsulated photoconverted HeLa mEos3.2 cell quantification (n = 4). F) Workflow of the sequential GRACE volumetric bioprinting and selective photostimulation process on microparticle embedded HeLa cells expressing mEos3.2. (i) View of the printing volume within the vial, showing the embedded alginate microspheres and their 3D reconstruction upon light sheet scanning. (ii) View of the feature-driven printed model (volume = 308 mm^3^) resulting in a channel network targeting the embedded alginate particles. (iii) View of the selective stimulation by delivering 405 nm light projections to the targeted microspheres. G) Volumetric printing shape fidelity quantification for negative features (channels diameter) and positive features (channels proximity to the alginate microsphere) (n = 5). H) GRACE ability to target and photostimulate encapsulated HeLa mEos3.2 cells, screening between spheres 600 μm apart. Scale bar: 500 µm. I) Photoconverted HeLa mEos3.2 quantification on targeted and non-targeted alginate microspheres (n = 4). J) Workflow of the sequential EmVBP and GRACE-mediated selective photostimulation on extruded features laden with HeLa cells expressing mEos3.2. (i) EmVBP + GRACE process consisting in embedded extrusion bioprinting and subsequent context-aware volumetric overprinting. (ii) After the washing step, coordinates registration and 3D reconstruction of the extruded features within the printing volume (324 mm^3^) is performed. (iii) Context-aware selective stimulation by delivering 405 nm light projections to the target extruded filament. K) GRACE enables to target and trigger photoconversion of mEos3.2 to a single filament, resulting in photoconverted HeLa cells. Scale bar: 1 mm. L) Close up photoconversion of the embedded extruded HeLa cells expressing mEos3.2 in either their native and photoconverted state in the targeted (i) and non-targeted (ii) extruded filaments. Scale bar: 1 mm. M) A higher percentage of photoconverted mESO3.2-expressing HeLa cells has observed in the stimulated filament (69.4 ± 6% of photoconverted HeLa cells) compared to the non-stimulated one (5.2 ± 0.6% of photoconverted HeLa cells) (n=4). Data represent mean ± SD. Significance was determined by one-way ANOVA (A) or unpaired t-test (I, M) *** = p <0.001.

As a proof-of-concept of selective microsphere targeting and photoconversion, encapsulated HeLa mEos3.2 cells were dispersed in the printing volume and scanned to register their 3D position (**Figure 4Fi**). A features-tuned (e.g. a perfusable channel network targeting microencapsulated HeLa cells) model was generated and volumetrically printed (**Figure 4Fii**). Next, the printed construct was placed back in the printer and re-scanned to detect its coordinates and orientation. The desired microparticles to be targeted for photoactivation were selected, resulting in the generation of light patterns specifically targeting the selected beads (**Figure 4Fiii**). More specifically, for the volumetric printing step, a printing shape fidelity of 88.3 ± 8% for the channels inner diameter and of 82.6 ± 5% for the channel proximity to the alginate microspheres was measured (**Figure 4G**), in line with the reported values in literature for volumetric printing shape fidelity.^[5,8]^ Results from **Figure 4H** showed how GRACE can precisely photostimulate encapsulated HeLa mEos3.2 cells, with the distinctive capacity to selectively tracking and targeting a single bead over the rotational motion of the vial, while also distinguishing the targeted bead from an adjacent sphere located within a 600 μm distance. Finally, a high photoconversion accuracy was also observed, resulting in 83 ± 4% of photoconverted cells in the targeted microspheres, compared to a 7 ± 2% in the adjacent non-targeted ones (**Figure 4I**), further confirming particle-selective activation. It should be noted that object recognition via the GRACE workflow could potentially be performed also with imaging modalities besides fluorescent imaging (i.e. bright field imaging for the spheroids), though we explicitly decided to use tagged cells, due to the improved contrast offered by light sheet fluorescence. After testing the ability of GRACE to target 3D specific features within a bioprinted construct, we investigated the applicability of this concept to model structures containing two cell-dense filaments produced via EmVP (**Figure 4J**). To maximize efficient on-target delivery, context-aware printing permitted editing the light projection calculations, to perform an obstacle-avoiding adjustment which reduced the light dose reaching the filament that was meant to remain non-stimulated (**Supplementary Figure S10**). From fluorescent microscopy pictures of the native and converted channels, we observed how the targeted filament qualitatively presented more photoconverted HeLa mEos3.2, compared to the untargeted one (**Figure 4K**). This difference was confirmed also by close-up images of the two filaments compared (**Figure 4L**), where significantly more photoconverted cells were observed in the stimulated target, resulting in a 69.4 ± 6% of photoconverted cells, compared to the non-stimulated (5.2 ± 0.6%) (**Figure 4M**). Overall, these results indicate that via context-aware printing, it is possible to pinpoint desired clusters of cells which can be followed and tracked also in real time as they rotate during the movement of the vial in the printer, enabling photo-stimulation not only by light patterning with static projections, but also through zonal volumetric light delivery.

### 2.5 Optogenetic control of gene expression via a NIR light-triggered synthetic circuit

While photoconvertible proteins like mEos3.2 are particularly relevant for applications in cell tracking over time, other class of light-sensitive proteins, optogenetic photoswitches, enable control of gene and protein expression. Optogenetic gene expression circuits typically comprise a sensing unit (often a protein, mammalian in origin or not), and the related downstream pathways leading to actuation. Several groups studied optogenetics to elegantly regulate gene or protein expression,^[58,59]^ to promote cell differentiation,^[60]^ and to tune mechanical processes,^[61]^ among other applications. Building on the findings reported in the previous paragraphs, we tested the feasibility of triggering gene expression via optogenetic circuits stimulated by the light projections of the volumetric bioprinter. To do this, we adapted a previously reported optogenetic gene switch,^[62]^ which has been originally engineered for firefly luciferase expression upon illumination with near-infrared light (740-750 nm). Briefly, this light-controlled switch is generated by fusing the bacterial phytochrome BphP1 and, its binding partner, QPAS1 to different units of specific transcription activator systems (e.g. Gal4/UAS). Excitation with near infrared light leads to QPAS1 and BphP1 to form heterodimers, bringing together the two units (Gal4 and VP16) necessary to trigger gene expression. In this study, as first proof of concept, we generated two gene transfer vectors that allow the optogenetic system to promote the expression of two transcription factors involved in the identity of pancreatic β-cells, *PDX1* and *MAFA*, and normally not expressed by HeLa cells, upon illumination with near infrared light (**Figure 5A**). The first gene transfer vector encoded the bacterial phytochromes (BphP1 and QPAS1) fused to the units necessary (Gal4 and VP16) to trigger specific gene expression. The second gene transfer vector encoded a inducible transcriptional reporter for the expression of *PDX1* and *MAFA* after near infrared light exposure. After transient transfection of these plasmids into HeLa, optogenetic responses were initially characterized via an 740 nm (NIR) LED-mediated illumination. Immunofluorescence staining showed how 24 h of pulsed NIR light exposure resulted in a higher induction of Pdx1 (yellow) and MafA (magenta) expression in 2D cultured samples (**Figure 5B**), and also in cells encapsulated in GelMA in 3D (**Figure 5C**). Firstly, in 2D culture, significantly increased levels of fluorescence per cell (2.9 ± 1.5 and 6.92 ± 4.62 fold-increment for Pdx1 and MafA respectively, **Figure 5D**), as well as higher number of cells expressing the target proteins (19.42 ± 12.81% unexposed vs 56.4 ± 2.22% illuminated, and 11.73 ± 6.6% unexposed vs. 37.62 ± 10.3% illuminated) for Pdx1 and MafA respectively, **Figure 5E**) were detected, when compared to non-illuminated controls. Co-induction of the two target transcription factor was also evidenced by co-localization in the nuclei analysed by Mander’s correlation index (0.97 ± 0.001 unexposed and 0.98 ± 0.01 illuminated) (**Figure 5F**), and increased gene expression upon NIR illumination was also confirmed via qPCR (Pdx1: 1.63 ± 1.95 dark vs 3.62 ± 1.89 illuminated, and MafA: 1.71 ± 2.1 dark vs 4.1 ± 2.57 illuminated) (**Figure 5G**). Over time after stimulation, the protein expression levels gradually return to baseline values (by 72-96 hours for Pdx1 and by 48-72 hours for the second target, MafA, **Supplementary Figure S12**). This is in accordance to the fact that this system relies on transient transfection, and that gene expression is conditional to light illumination, and therefore better suited to study the effect of temporary alteration of protein synthesis. A first limitation of the system was also revealed, as only a fraction of the cells could be successfully transfected, and the amount of NIR-light responsive cells appeared to be lower in 3D samples, compared to 2D culture. Of note, transfected cells displayed lower viability values after encapsulation, with ⁓70% of living cells after 1 and 3 days, compared to values of ⁓88 and 82% for the untreated HeLa (**Supplementary Figure S13)**. Moreover, some degree of baseline activation was still observed in samples left in the dark, a common challenge in optogenetics. Regarding the 3D culture within GelMA hydrogels, the response to optogenetic stimulation was further assessed as a function of the delivered light dose, showing how the intensity of the immunofluorescence signal, proportional to the amount of expressed protein in response of the light stimulation, sharply increased to a plateau 2.12 fold (Pdx1), and 8.92-fold (MafA) for light intensity values of 0.1 mW/cm^2^ and higher, when compared to the baseline value for non-illuminated samples (**Supplementary Figure S14**). Partial activation was also observed for intensities as low as 0.05 mW/cm^2^, suggesting that tight control of the delivered light doses is necessary for providing high contrast between in-target and off-target (non-stimulated) regions of the hydrogels.

**Figure 5.**
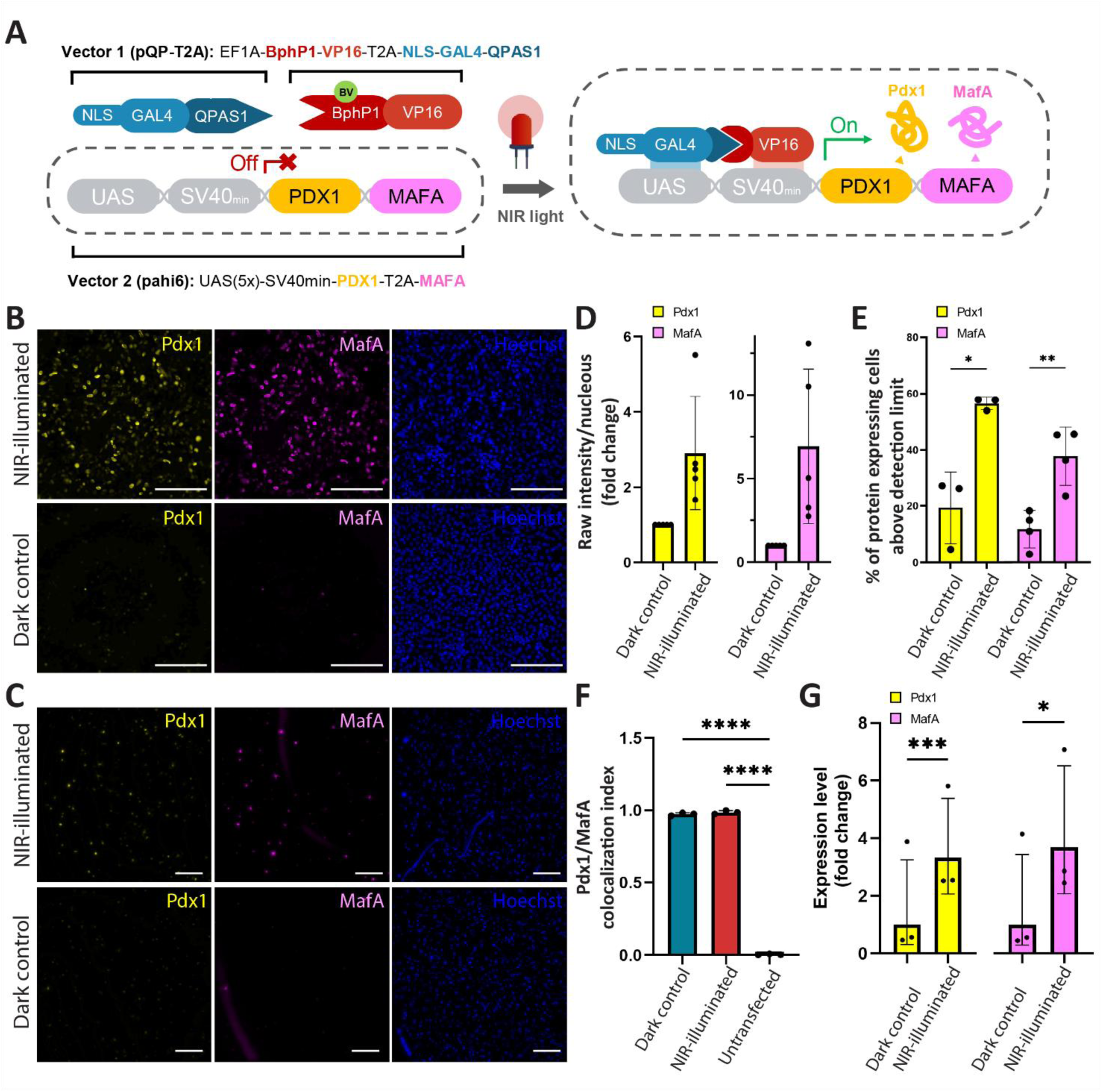
Near-infrared (NIR) light-triggered expression of Pdx1 and MafA in 2D and 3D HeLa cell culture. A) Schematic representation of QPAS1-BphP1 optogenetic induced expression of *PDX1* and *MAFA*. B) Immunofluorescence characterization of 2D-cultured HeLa cells transfected with optogenetic vectors 1 (pQP-T2A) and 2 (pahi6) after 24 h of pulsed (30 s ON, 180 s OFF) NIR illumination using a 740 nm LED at 25 mW/cm^2^ leads to increased Pdx1 (yellow) and MafA (magenta) expression compared to the transfected HeLa cells maintained in the dark as control. Cell nuclei were stained with Hoechst (blue). Scale bar: 200 µm. C) Immunofluorescence characterization of 3D-encapsulated HeLa cells (2 × 10^6^ cells/mL) transfected with optogenetic vectors 1 (pQP-T2A) and 2 (pahi6) after 24 h of pulsed (30 s ON, 180 s OFF) NIR illumination using a 740 nm LED at 25 mW/cm^2^ leads to increased Pdx1 (yellow) and MafA (magenta) expression compared to the transfected HeLa cells maintained i the dark as control. Cell nuclei were stained with Hoechst (blue). Scale bar: 200 µm. D) Quantification of total raw image fluorescent intensity of Pdx1 (yellow) and MafA (magenta) normalized to number of nuclei in 2D HeLa cultures after 24 h of pulsed light illumination. Values are expressed as fold change relative to the corresponding dark control. E) Percentage of Pdx1 (yellow) and MafA (magenta)-expressing cells in 2D after 24 h of pulsed light illumination. F) Pdx1 and MafA signal correlated in the fluorescence measurements confirmed by a Meanders’ correlation coefficient. G) *PDX1* (yellow) and *MAFA* (magenta) RNA expression after 24 h of near-infrared pulsed illumination, expressed as fold change relative to the average of unilluminated controls and normalized to *GAPDH* housekeeping gene. Data represent mean ± SD. Significance was determined by unpaired t-test (E), one-way ANOVA (F) or paired t-test (G). * = p< 0.05, ** = p< 0.01, *** = p< 0.001, **** = p< 0.0001.

### 2.6 Zonal optogenetic gene transcription induction in volumetrically bioprinted hydrogels

After assessing the possibility to induce gene expression in transfected HeLa cells in 2D and in 3D, we tested if the optogenetic triggering was possible in volumetric bioprinted structures using the tomographic bioprinter as light source. While the light-induced heterodimerization of the photoswitches (QPAS1 and BphP1) occurs in seconds to minutes, gene expression, and particularly its downstream protein production requires timeframes in the order of hours, to meaningfully influence cell behavior. To enable experiments over multiple hours in cell friendly conditions, we adapted the printing set-up with a custom-made temperature control unit, in which the refractive index matching bath in which the bioresin-holder vial is typically immersed, is heated to 37°C (**Figure 6A**) Gas exchange was ensured by a filter cap, and pH was regulated with HEPES in the culture media. These temperature levels are crucial to ensure optimal cell bioactivity for the whole duration of the optogenetic activation experiment (20 h), where a thin band of 500 μm was projected with NIR 750 nm wavelength laser above the sample base. First, a viability assay performed after 20 h of NIR light shining, under the same 30s ON - 180s OFF pulsed regime (at 0.6 mW/cm^2^), showed a high viability throughout the whole construct volume (**Figure 6B**), proving the safety of both the volumetric printing step and the prolonged, intermittent stimulation with NIR laser. After a 20 h spatial-selective illumination, we observed significantly higher Pdx1 and MafA expression with zonal localization corresponding to the shone region, when compared to the non-illuminated regions of the hydrogel construct (**Figure 6C**). Nevertheless, a diffuse pattern of activation was also shown in segments adjacent to the target (up to approximately 750 µm away from the border of the light projection), and decreased further away from the illuminated region, reaching baseline levels (**Figure 6D**). This is likely due to light scattering within the bioprinted gel, with scattered light delivering a sufficient dose to partially activate cells, therefore reducing spatial resolution proportionally to the cell concentration within the construct (**Supplementary Figure S15**). Modelling the light scattering profile is therefore especially important, as its effects can be predicted and mitigated also by modifying the pattern of the projected light, as well as tuning the delivered light intensity to limit off-target stimulation. Overall, in our experiments, activation with the NIR light projections delivered by the printer resulted in higher induction of Pdx1 and MafA, with respectively 3.62 ± 2.04 and 8.13 ± 4.75 fold increment in fluorescence intensity in the nuclei over the unstimulated baseline (**Figure 6E**). Consistently, the percentage of successfully activated cells significantly increased from 6.51 ± 0.54 and 3.89 ± 2.93 (baseline levels for Pdx1 and MafA) to 35.68 ± 10.59 and 33 ± 10.68% in the on-target region of the hydrogel (**Figure 6E**). Additional pilot tests to confine the activation, within projected patterns generated by the printer were also conducted in cell-laden GelMA hydrogels, showing preferential activation within the illuminated regions, which resulted in a 6.28 ± 2.58 (Pdx1) and 5.12 ± 0.79 (MafA) fold increase in the density of activated cells within star-shaped pattern (**Supplementary Figure S16)**. Albeit preliminary, these results open the possibility to create hydrogel systems in which gene and protein expression levels in the bioprinted cells can be altered locally, to better study how varying patterns of expression influence tissue behaviour. Further optimization of this optogenetic activation circuitry should however be warranted, to unlock the capability to fully perform targeted tomographic activation between a volume of hydrogel, in a similar fashion to what we showed when using the faster reacting mEos3.2 photosensitive proteins. To address the limitations identified in our work, additional optimization steps are thus advantageous to maximize resolution, and contrast over off-target areas, to enable the production of more complex light-induced patterns. In particular, the use of engineered cell lines that stably express the synthetic circuits would be preferrable over the transient expression used in our study, which is inherently limited by transfection efficiency. This would be of particular importance when using especially sensitive cell types, such as primary adult stem cells or pluripotent stem cells,^[60]^ which are notoriously difficult to transfect with high efficiency. Additionally, strategies such as refractive index matching within the bioresin,^[42]^ and computational corrections of the light projections,^[54,63]^ could be explored to reduce scattering-mediated artefacts, and therefore also assessing the potential for printing and photo-stimulating at higher cell densities. Furthermore, specific to the QPAS system, the circuit could be inactivated with red light, thus, combining different wavelength to activate or inhibit the reaction could be a strategy to improve contrast.^[64]^ Moreover, while in this study we used a well-established NIR-responsive system, more efficient circuits (reaching up to 135-fold improved gene expression upon illumination) have been designed more recently.^[65]^ Likewise, different designs relying on light-responsive CRISPR-based gene editing and gene activation systems, could also be of great interest to reduce the timeframe needed for photostimulation.^[66]^ The implementation of more advanced synthetic circuits could help improving signal-to-noise ratio, while also limiting the risk of partial circuit activation in non-illuminated conditions, permitting to unlock the full potential of our approach. Moreover, in future studies repeated photostimulation steps could be performed at different time points during culture, simply moving the sample back in a tissue culture incubator when the light exposure is not required. Alternatively, longer illumination timeframes could be also envisioned, though the addition of a gas exchange system to more tightly control pH and oxygen tension in the printing chamber should be envisioned. Overall, our results provide the first demonstration and proof-of-concept that spatial control of optogenetic responses with light provided by a volumetric bioprinter is feasible, opening promising avenues for further research and for the development of engineered tissue models with remotely controllable cell behaviour.

**Figure 6.**
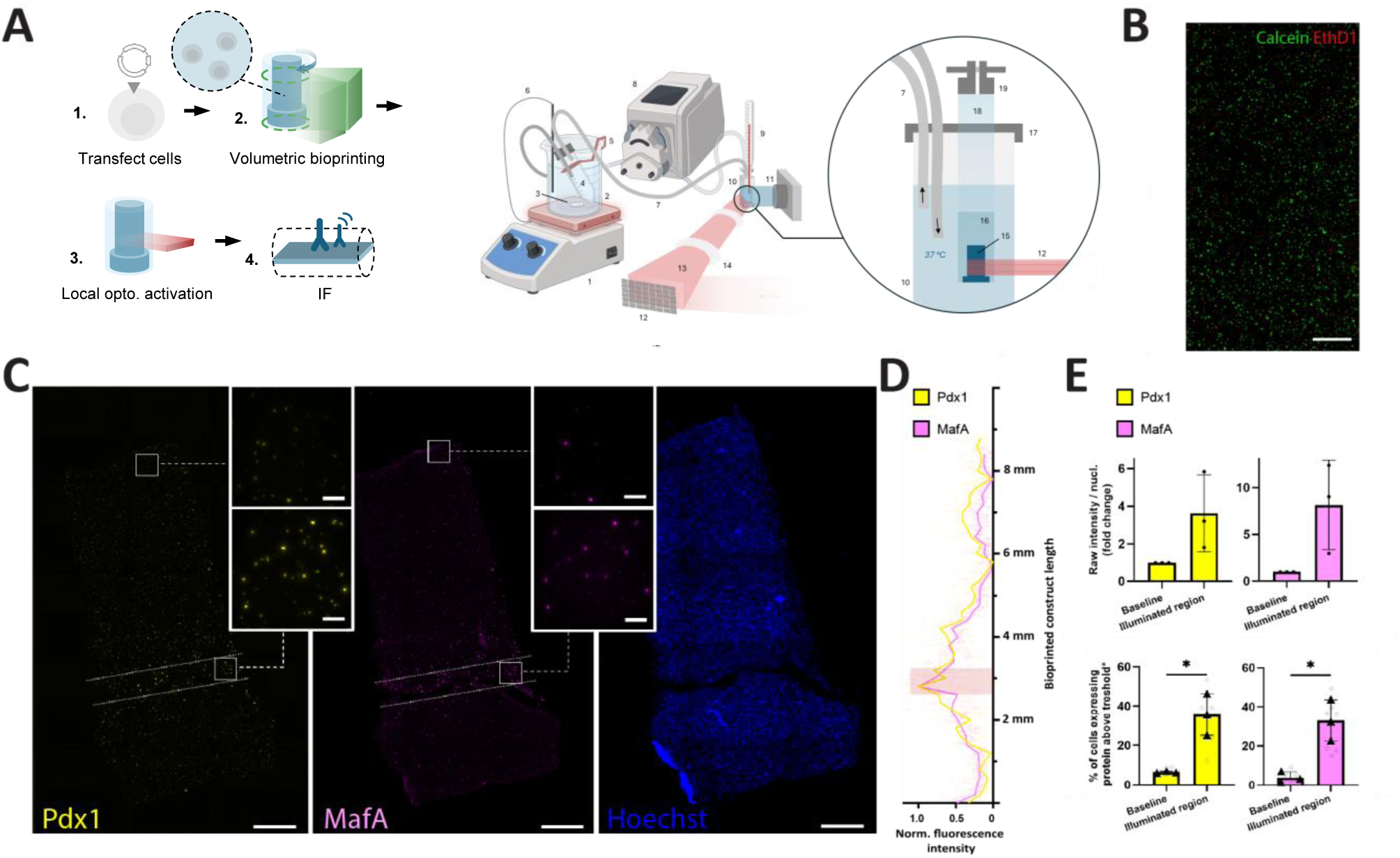
Spatially-selective optogenetic-induced expression of Pdx1 and MafA using light projections of a volumetric printer. (A) (Left) Schematic of workflow for the preparation, bioprinting, spatially-selective illumination and immunofluorescence (IF) characterization of cell-laden bioprinted constructs. (Right) Schematic representation of the setup for spatial optogenetic induction of Pdx1 and MafA expression using a volumetric bioprinting. A closed-loop perfusion system was used to maintain bioprinted constructs at 37°C during 20 h optogenetic stimulation. A heating plate (1) warms a beaker (2) filled with water and equipped with a magnetic stirring bar (3) to ensure uniform temperature distribution. A sealed 50 mL tube filled with demi water (4) and connected to in- and outflow tubing is held submerged by a custom 3D-printed holder (5). Temperature is monitored by a temperature probe (6) placed inside the beaker. Warmed demineralized water is circulated via tubing (7) through a peristaltic pump (8) into a cuvette (10), maintaining the thermal environment around a bioprinted, cell-laden construct (15) embedded in phenol red free-medium supplemented with 25 µM biliverdin. The construct is cultured inside a printing vial (18), positioned trough an opening in a 3D-printed cap (17) containing tubing inlets and sealed with a sterile filter to ensure oxygen exchange (19). Laser projections (12) are directed via lenses (14) and focused onto the vial using a laser beam (13) for targeted photoactivation. An orthogonal camera (11) assists in real-time projection alignment. Image uses drawings from Biorender. (B) Live (calcein)/dead (ethidium homodimer-1) assessment of transfected HeLa cells in volumetrically printed GelMA hydrogels, after 24 h of optogenetic induction in the volumetric bioprinter. (C) Immunofluorescence characterization of the spatial-selective optogenetic expression of Pdx1/MafA in HeLa-laden (2.0 × 10^6^ /mL) volumetrically printed GelMA constructs (volume = 255 mm^3^) using static projections from a volumetric printer. A light band of 0.5 mm thickness (λ=750 nm) is projected into a bioprinted cylindrical construct laden with HeLa cells at a light intensity of 0.6 mW/cm^2^ during 20 h. (D) Assessment of the spatial selectivity of gene expression. Graph represent normalized pixel intensity side-profiles of previous IF staining. Red boxes mark the projection area. (E) Quantification of raw image fluorescent intensity normalized to number of nuclei within dark and stimulated regions of the HeLa-laden constructs. Values are expressed as fold change relative to the corresponding dark control (N = 3, n = 3). Detectable number of cells expressing Pdx1/MafA as percentage of total nuclei after image thresholding in dark and stimulated regions of the construct. Triangles represent biological replicates, grey dots technical replicates (N = 3, n = 3). Scale bar in B is 1 mm. Scale bar in C is 1 mm, inset is 200 µm. Data represent mean ± SD. Significance was determined by unpaired t-test, * = p < 0.05.

## CONCLUSIONS

In this study, we demonstrated a novel strategy to integrate spatiotemporal control of cellular events into bioprinted constructs, by extending tomographic volumetric bioprinting functions beyond the conventional material crosslinking. Leveraging light tomographic projections (405 nm and 750 nm), we showed that spatially selective activation of photosensitive proteins and optogenetic circuits can be achieved post-printing within centimeter-scale hydrogel constructs, with a resolution of 250 µm. Rapid, cytocompatible region-specific activation of photoconvertible proteins (mEos3.2) was shown via tomographic light dose delivery, with high photoconversion efficiency (>80% of the cells in the targeted areas). This approach enables a new level of control over cellular processes in a 3D environment, solving potential limitations of conventional stimulation methods which either lack penetration depth (i.e. two-photon stimulation) or spatial specificity in 3D (such is the case of photomasks). Furthermore, by integrating this approach with imaging and computer vision-based feedback strategies, we show the feasibility of context-aware stimulation, where light can be selectively delivered to features or regions of interest that are automatically identified and registered by the printer itself, with minimal input from the user. In future developments, the GRACE workflow could also be applied to facilitate registration of the shape, orientation and positioning of constructs that undergo shape changes during culture, in order to identify and photostimulate regions of interest within the construct. Finally, the proof-of-concept of activating NIR light-controlled optogenetic gene expression circuits using a multi-wavelength volumetric bioprinter opens exciting future opportunities for a minimally invasive modulation of biological functions within thick and complex biofabricated models. Challenges in achieving high transfection efficiency, and low signal-to-baseline contrast we encountered for this specific NIR system, were identified, and indicate directions for improving the technique. Looking ahead, this technology could be extended to control multiple cell types, by leveraging different wavelengths and optogenetic circuits, to dynamically modulate cellular interactions over time, particularly when combined with advances in synthetic biology and programmable materials. Overall, the convergence of optogenetics and multi-technology volumetric bioprinting represents an encouraging step toward smart biofabrication processes, shifting from statically cultured constructs to stimulation systems that can precisely control cellular processes in space and time.

## EXPERIMENTAL SECTION

### HeLa mEos3.2 generation

HeLa Flp-in cells stably expressing a TetR and carrying a single genomic FRT site were cultured in DMEM high glucose medium supplemented with 9% fetal bovine serum and 1% penicillin/streptomycin at 37°C with 5% CO_2_. All cell lines were routinely screened (every 8–12 weeks) to ensure they were free from mycoplasma contamination. Cloning pCDNA5-H2B-mEos3.2, a mammalian expression vector for the human histone H2B tagged with the green/red photoconvertible fluorescent protein mEos3.2, was derived from pH2B-mIRFP670nano3 (Addgene plasmid #127438;http://n2t.net/addgene:127438; RRID:Addgene_127438), and pCDNA5/FRT/TO (Invitrogen) by conventional molecular cloning. The HeLa-H2B-mEos3.2 stable line was generated using flp/frt recombinase-mediated cassette exchange. Maternal HeLa Flp-in T-Rex were transfected with the donor plasmid pCDNA5-H2B-mEos3.2 and pOG44 (Invitrogen), carrying Flp recombinase. Isogenic cells stably expressing H2B-mEos3.2 from a doxycycline-sensitive promoter were selected by treatment with 200 μg/ml hygromycin B (InvivoGen, ant-hg-5). At least 24 hours prior to the start of any of the experiment, the culture media was supplemented with doxycycline (1 µg/mL), to ensure expression of the mEos3.2 protein.

### HeLa mEos3.2 photoswitch light dose threshold test with 405nm microscope laser

2D plated, doxycycline-treated HeLa mEos3.2 cells were exposed to a range of light intensities and overall doses using the 405 nm laser of a Thunder microscope (Leica). Cells were irradiated using a 10x magnification lens, at different laser intensities (75.8, 160.2, 243.4 and 323.6 mW/cm^2^) and doses (2.4, 4.8, 7.2, 9.6, and 14.5 J/cm^2^) and imaged before and after irradiation (n=3). Cell photoconversion percentage was analysed in FIJI. Cell ROIs were identified in the pre-stimulation cell channel (Ch1) by Otsu thresholding (dark background), watershed separation of touching cells, and particle detection (minimum area 25 px²). For each ROI, activation was assessed in the mEos3.2 reporter channel (Ch2) as the median of the pixel-wise (post − pre) intensity difference within the cell footprint, normalised to the median pre-stimulation intensity of the same cell. A cell was classified as activated when this relative change exceeded 30%.

### HeLa mEos3.2 photoswitch light dose threshold test with the volumetric printer

To characterise the activation behaviour of the mEos3.2 system in 3D, HeLa mEos3.2 were casted at a concentration of 2×10^6^ cells/mL in 1.5 mm-thick GelMA constructs and treated with doxycycline overnight. Subsequently, the constructs were placed back in the printer in a squared cuvette and exposed to 405 nm light projections of a series of 2D circles (500 µm in diameter) under various laser intensity (20 – 50 mW cm^−2^) and dose (1000 – 20000 mJ cm^−2^). Samples were imaged with a Thunder microscope and mean grey values at the irradiated regions obtained using FIJI.

### Volumetric bioprinting of cell-laden hydrogels

The pivotal principle in volumetric printing is the threshold-based photochemical behaviour that initiates the material crosslinking. Briefly, tomographic volumetric bioprinting consists in delivering visible 2D light projections to a rotating vial containing a photoresponsive, cell-laden hydrogel, using a spatial light modulator (i.e., a digital micromirror device, DMD). The projections corresponding to each angle of rotation of the vat are calculated following a tomographic reconstruction algorithm,^[46,47,67]^ resulting in a cumulative light dose that exceeds the crosslinking threshold of the biomaterial only in the voxels corresponding to the object to be printed, solidifying the whole structure at once and in a layer-free fashion. In this study, a Tomolite v2.0 (Performance version) volumetric bioprinter (Readily3D SA, Switzerland) and a Tomolite v2.0 (open-format version) volumetric bioprinter with a multi-wavelength add-on (Readily3D SA, Switzerland) were used to generate the 3D constructs. The GelMA (degree of methacrylation = 60-80%, molecular weight = 90kDa, Rousselot, Belgium) and AlgMA (TNT-05, degree of methacrylation = 24%, molecular weight = 250-350kDa, Polbionica, Poland) resins were dissolved at concentrations of 5% w/v and 3% w/v in a PBS-LAP stock-solution (1x PBS, 0.1 mg/mL LAP). The GelMA bioresins (cell-free or embedding 1-2×10^6^ cells/mL) were poured at 37°C in cylindrical glass vials (diameter: 10 mm) and kept at 4°C to ensure thermal gelation. User-designed CAD files were loaded and processed using the Readily3D Apparite software. A 9.9 mW/cm^2^ average light intensity was set and a light dose of 180 – 230 mJ/cm^2^ and 372 mJ/cm^2^ was used to crosslink the GelMA and AlgMA bioresins, respectively. After printing, the vials were washed with pre-warmed PBS at 37°C to remove the uncured bioresin from the printed structure. The tomographic activation of a cylinder inside a printed disk (d=1mm, volume = 4.71 mm^3^) was calculated with the Object-Space Optimization algorithm,^[68]^ using D_l_ = 0.7 and D_h_ = 0.9 after 40 iterations, using a 180° projection strategy, taking advantage of the π symmetry for tomographic reconstructions. The target cylinder (6.4mm diameter, 6mm height, 2×10^6^ cells/mL HELA mEos3.2) was irradiated with a mean in-part voxel dose of 10.7 J/cm². The choice of the cylindrical sample was made to facilitate imaging the activated slice, when cutting the sample transversally. To challenge the system, the region to be stimulated was placed on purpose away from the center of the disk, so that the light patterns to be delivered by the printer change with the angle of rotation, differently than what would have happened had the cylindrical region been centered exactly on the rotation axis of the vial.

### Spatial activation fidelity and volumetric printing shape fidelity

Both the activation and printing shape fidelity (%) indexes were calculated as reported in Equation 1,^[69]^ where “m” and “M” refer to the sample measured feature and the known designed feature dimension, respectively.

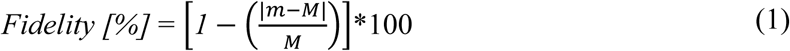

### Rheology on low MW GelMA fluid bulk support bath for embedded extrusion printing

The 90 kDa GelMA bioresin, meant to be used as a photocrosslinkable support bath, was dissolved at concentrations of 5% w/v in a PBS-LAP stock-solution (1x PBS, 0.1 mg/mL LAP) at 45°C. The rheometer (HR-2, TA Instruments, New Castle, DE, USA) testing temperature was set to 21°C. The shear rate ramp test was performed with a frequency of 1 Hz and shear strain of 1%. The shear stress and the viscosity were measured during a logarithmic shear rate ramp from 0.01 to 1000 s−1 (2 seconds per point). The shear stress ramp test was similarly performed, now using the shear strain during a logarithmic shear stress ramp from 0.01 to 1000 Pa (2 s per point). The stress recovery of the GelMA or gelatin was performed using a thixotropy test: similarly, the temperature to 21°C (tolerance 0.1°C for 1 min to allow temperature equilibration) and the G’ and G’’ were measured for 2 min with a frequency of 1 Hz and shear strain of 1% (2 s per point). Then a high shear rate was applied (500 s−1) to disrupt the gel for 2 min (2 seconds per point). After, the G’ and G’’ were again measured for 10 min, with a frequency of 1 Hz and shear strain of 1% (2 s per point).

### Embedded Extrusion (bio)printing

3D (bio)printing experiments were performed with a R-GEN 100 (RegenHu, Switzerland), via a pneumatic-driven extrusion printhead. A fluid bulk support bath, developed in a precedent study, was used. Briefly, a low molecular weight (MW = 90 kDa) GelMA (degree of modification: 80%) bioresin is prepared at 4% w/v concentration with 0.1% w/v LAP as initiator and used straightforwardly as support bath upon thermal gelation. Thanks to a smaller length of the polymer chains, this formulation showed self-healing-like properties, preventing the needle from creating grooves or scratches in the bath during the translation movement, thus affecting the bioink deposition quality.^[7]^ For evaluating the printing resolution, 3 mL syringes were loaded with 1 mL of Cy-3.5 stained 2% w/v methylcellulose bioink (already characterized in a previous work).^[6]^ A 27G needle, two different printing pressures (10 and 20 kPa), and four different printing speeds (0.5, 1, and 1.5 mm/s) were tested to characterize the filaments’ width. Images (n = 6) of the embedded extruded filaments were taken with a microscope (Thunder microscope, Leica Microsystems, The Netherlands) and analyzed with ImageJ. For HeLa mEos3.2 cells bioprinting, cells were mixed with the methylcellulose at a 30 × 10^6^ cells/mL concentration and extruded with a 20G needle.

### Embedded Extrusion Volumetric Printing

The extruded components and the geometries meant to be volumetrically overprinted were designed and saved as independent STL files. The GelMA for the preparation of the fluid bulk support bath was dissolved at 4% w/v concentration in a PBS-LAP stock-solution (1x PBS, 0.1 mg/mL LAP) at 37°C and left to thermally gelate overnight at room temperature. A printing vial was loaded with the fluid GelMA resin, to serve as support bath, and placed in a R-GEN 100 extrusion printer (RegenHu, Switzerland). A 3 mL syringes were loaded with 1 mL of bioink (2% w/v methylcellulose + 30×10^6^ HeLa mEos3.2 cells/mL), and the G-codes of the extruded features were manually written. Subsequently, the vial was loaded into a custom-made multi-wavelength Tomolite volumetric printer (Readily3D SA, Switzerland). To align the features extruded in the previous step and the model for volumetric overprinted, a first series of light-projections was delivered on the rotating vial with a 520 nm laser, to prevent unwanted cross-linking. Therefore, volumetric bioprinting is performed as described above. At the end of the EmVP process, the final construct was anchored at the base to the printing vial. Finally, the vial was heated to 37°C to dissolve the unpolymerized support bath and the construct was retrieved and washed with pre-warmed PBS.

### Zonal HeLa mEos3.2 photoswitch stimulation within printed constructs generated with volumetric bioprinting or EmVP

A first volumetric or extrusion-volumetric print of the desired model is performed as described above. The unpolymerized resin is washed out, and the vial containing the structure printed with the first material is then filled with PBS. To prevent the displacement of the first print during the washing steps, a base, made with the same bioresin of the construct, was printed at the bottom of the vial, to anchor the printed object in place. For light-stimulation experiments, a pre-designed photomask was loaded in the Polychromatic Tomolite volumetric printer. The samples and the projection pattern were aligned manually, using a 520 nm wavelength light source while avoiding any unwanted photoconversion of HeLa mEos3.2 cells. Finally, the light pattern was projected at 405 nm onto the anchored structure at a set light dose value. The anchoring base was ultimately removed at the end of the process, by cutting it with a razor blade.

### Alginate microspheres production and microencapsulation of HeLA mEos3.2

An extrusion printer (Inkredible, Cellink Sweden) was used as air pressure regulator, by connecting two independent printheads connectors to a coaxial needle (r=0.3/0.69mm l=8mm, RegenHU, Switzerland). More specifically, the inner needle was used for bioink extrusion and the outer one for the air flow to break the bioink filament into droplets.^[70,71]^ Sodium alginate (71238, Sigma-Aldrich) was dissolved at a 2% w/v concentration (eventually mixed with HeLa mEos3.2 cells at 30×10^6^ cells/mL concentration for experiments involving living cells) and loaded into a 3 mL cartridge to be connected to the air-assisted coaxial nozzle extrusion system. For the characterization of the achievable microspheres size, the air pressure was set between 30 and 120 kPa, and the alginate extrusion pressure was kept between 20 - 30 kPa. The microspheres were collected into a beaker containing a crosslinking solution of 50 mM CaCl₂ and kept there for a maximum of 3 minutes. The microgels were finally collected with a cell strainer and thoroughly washed with PBS.

### GRACE and particle tracking for selective protein photoactivation on encapsulated HeLa mEos3.2 cells

Vybrant™ DiD (ThermoFischer Scientific) was used as cell-labelling membrane staining on HeLa mEos3.2 for the detection step within the GRACE process. For the GRACE procedure and subsequent photoswitch activation, a custom-built volumetric printer was used.^[11]^ Cell-laden alginate microparticles were produced as described in the previous section, resulting in beads of 1.18 ± 0.13 mm in diameter, and with a cell load of 30×10^6^ cells/mL, accounting for an estimate of approximately 27000 cells/bead, assuming the volume of a single bead is that of a sphere having the average microparticle diameter. The cell-laden alginate microparticles were suspended in a 10% w/v GelMA resin and loaded into the printing vial, gently suspending them by agitating the volume while the mixture was kept in an ice bath until the GelMA resin thermally gelated. The volume was then scanned with 650 nm light over Θ_total_= π, and the centroid coordinates of each alginate sphere was calculated by a cluster detection algorithm as previously described.^[11]^ Following the GRACE workflow, the adaptive models, consisting of a set of channel networks, were parametrically generated by the software to target the embedded alginate beads and sculpted in the GelMA volume using tomographic volumetric printing. Channels were programmed to graze the beads at a distance of 300 μm and to display an inner diameter of 800 μm. Subsequently, the printed constructs were retrieved, washed to remove all the uncured material, and placed back in the volumetric printing vial while resuspended in 5% w/v gelatin, able to keep the construct in a fixed position within the volume upon thermal gelation. A second scan was performed with NIR light to register the new microspheres coordinates. Therefore, upon specific selection of the microspheres to stimulate, new adaptive models, consisting in spheres embedding the whole volume of the targeted alginate microbead, were generated and shined with 405 nm according to the GRACE workflow. To prevent the laser to collimate on possible obstacles (different alginate microbeads) in the shining trajectory during the stimulation step, the 2D light projections were generated including a shadow correction step, crucial to turn the DMD off when an obstacle was interposed between the light source and the targeted microbeads. After the stimulation step, the gelatin holding the bioprinted construct in place was dissolved and the model retrieved for further analysis.

### GRACE for selective protein photoswitch of HeLa mEos3.2 cells within printed constructs generated with EmVP

Vybrant™ DiD (ThermoFischer Scientific) was used as cell-labelling membrane staining on HeLa mEos3.2 for the detection step within the GRACE process. Two filaments spanning a length of 7 mm each embedding HeLa mEos3.2 cells (20 × 10^6^ cells/mL, mixed with 2% methylcellulose as carrier ink) were extrusion bioprinted into the previously described low MW GelMA bath, following the EmVP workflow described above. The filaments were then encased in a volumetrically printed prism. A tomographic illumination pattern was produced to stimulate the HeLA mEos3.2 cells in only one of the extruded filaments, which was mapped and targeted following the GRACE algorithm.

### Establishment of a NIR-responsive optogenetic circuit for triggering gene expression

Cells were engineered with a synthetic construct based on the *Rhodopseudomonas* phytochrome Bphp/QPAS1 system. As model proof-of-concept, the construct was engineered to trigger the contextual expression of two master transcription factors in pancreatic β-cell development, pancreatic and duodenal homeobox 1 (Pdx1) and MAF BZIP transcription factor A (MafA). The pSUBCMV plasmid encoding the Bphp/QPAS1 optogenetic construct for light-induced gene expression (*CMV-BphP1-VP16-T2A-NLS-GAL4DBD-QPAS1-pA*) was obtained from Addgene #102583,^[57]^ expanded in DH5a cells, and isolated using a NucleoBond Xtra Maxi kit (Macherey-Nagel) following the manufacturer’s instructions. *PDX1* and *MAFA* were cloned into a pcDNA backbone under a 5x(UAS) GAL4 binding site and a SV40min promoter (*5UAS-SV40min-PDX1-T2A-MAFA-pA*). The cloned construct was also expanded in DH5a cells and isolated using a NucleoBond Xtra Maxi kit (Macherey-Nagel) following the manufacturer’s instructions.

### HeLa cell culture and transient transfection of optogenetic switch

HeLa cells were plated in 6-well plates at a cell density of 600,000-800,000 cells/well. The day after plating, the medium was refreshed, and HeLa cells were transfected with both plasmid constructs at a 1:1 molar ratio using lipofectamine 2000 (ThermoFisher Scientific) following the manufacturer’s instructions. 6 h after transfection medium was supplemented with 25 μM biliverdin and the samples were left in the dark overnight. The day after transfection, the medium was refreshed to phenol-red free DMEM supplemented with 25 μM biliverdin. Cells were then either directly exposed to light regimes in 2D or further used for bioprinting experiments.

### Optogenetic induction of Pdx1 and Mafa expression in HeLa cells using LED illumination

2D transfected HeLa cells were exposed to pulsed light regimes of 30 s ON / 180 s OFF using 740 nm LEDs (Roithner, LED740-01AU), max intensity = 25 mW/cm^2^, connected to an Arduino Mega (Arduino) in phenol red-free cell culture media supplemented with 25 μM biliverdin. For the optogenetic induction within 3D constructs HeLa cells were bioprinted into 2×8 mm disks using a volumetric bioprinter at a cell density of 2 × 10^6^ cells/mL. Bioprinted disks were placed in phenol red-free cell culture media containing 25 μM biliverdin and bioprinted constructs were exposed to pulsed light regimes of 30 s ON / 180 s OFF using 740 nm LEDs (Roithner, LED740-01AU), 10 mW/cm^2^, connected to an Arduino Mega (Arduino). A 3D-printed support was used to both mount the cell culture plate and house the electronics for driving the illumination system from the bottom of the cell culture plate. Cells were analysed 24h after induction of light exposure via immunofluorescence, as described in the following sections. The light stimulation regimen utilized in this study was adapted from previous reports.^[62,64]^

### Zonal optogenetic induction of Pdx1 and MafA expression in volumetrically bioprinted constructs

The bioprinting and 3D light pattern exposure steps were adapted from the same protocol described for the zonal protein photoswitch experiments. Subsequently to the volumetric bioprinting step and anchoring of the construct to the vial, phenol-red free cell culture medium supplemented with 25 μM biliverdin and 25 mM HEPES was added in the vial instead of PBS, and the vial was finally sealed with a custom-made cap equipped with a sterile filter. Considering the long exposure time required for optogenetic gene activation (20 h), and to prevent zonal mistargeting caused by the shrinking of the construct occurring within 2-3 h after printing, the bioprinted constructs containing the transfected HeLa cells were first kept in the incubator at 37°C within the filter-capped printing vial, prior to performing the optogenetic stimulation experiments. The vials were therefore placed in the Polychromatic Tomolite volumetric printer (Readily 3D, Lausanne, Switzerland) where a custom heating stage was designed to assure temperature control of the refractive index matching liquid throughout the whole NIR light (750 nm) exposure step. More specifically, the refractive index matching liquid-containing cuvette was equipped with a 3D printed cap holding the tubing system of a peristaltic pump perfusion set up. This was connected to a closed loop perfusion system consisting of a falcon tube reservoir containing the warm refractive index matching liquid, submerged into heated up-distilled water. Finally the cuvette was connected back to the reservoir. A flow rate of 5 mL/min was set to keep the refractive index matching liquid around the printing vial at 37°C, and the temperature was monitored with a probe thermometer. Throughout the whole illumination time, the bioprinted construct was kept in the vial containing phenol red-free media with 25 μM biliverdin and 25 mM HEPES, 750 nm light was projected with 0.6 mW/cm^2^ intensity. The samples were illuminated with a pulsed-light regime of 30 s ON and 180 s OFF for 20 h, after which the samples were retrieved and stained for Pdx1 and MafA to assess the optogenetic activation. The regions of the bioprinted construct that were not illuminated were used as internal control. An additional bioprinted cylinder was kept in the dark in a conventional cell culture incubator as dark control.

### Immunofluorescence stainings of 2D cell culture

Following optogenetic activation 2D-cultured cells were fixed in 4% formaldehyde solution for 15 min at room temperature and permeabilized with 0.1% (v/v) Triton X-100 in phosphate buffered saline for 10 min. After permeabilization, cells were blocked with a blocking buffer (1% (w/v) BSA solution in PBS) for 1 h at room temperature. The cells were incubated overnight at 4°C with primary antibodies goat polyclonal anti-Pdx1 (1:300 dilution, R&D systems) and rabbit monoclonal anti-MafA [D2Z6N] (1:800, Cell Signaling Technology) diluted in blocking buffer. After primary antibody incubation the cells were washed three times with blocking buffer and then incubated with secondary antibodies donkey anti-goat AlexaFluor 488 (1:300 dilution, Thermo Fisher Scientific), donkey anti-rabbit AlexaFluor 555 (1:300 dilution, Thermo Fisher Scientific), and 1 μg/mL Hoechst 33342 (Thermo Fisher Scientific) diluted in blocking buffer for 1 h at room temperature in the dark. Samples were then washed three times in PBS and analysed on a Leica Thunder inverted microscope. Signal quantification was performed in Fiji (ImageJ) by threshold-based segmentation of hoechst-stained nuclei followed by watershed separation to determine nuclei counts per image. Mean fluorescence intensities for Pdx1 and MafA channels were background-corrected using user-defined ROIs and subtraction from whole-image means. These corrected values were normalized to nuclei counts to compute average per-cell fluorescence.

### Immunofluorescence stainings of cell-laden bioprinted constructs

Bioprinted constructs illuminated either with the LED system or the volumetric bioprinter itself were fixed in 4% formaldehyde solution for 45 min. For the Bioprinted constructs for which optogenetic gene expression was triggered with volumetric bioprinter itself, constructs were embedded in 5% agarose and sectioned into 400 μm sections using a VT100 vibratome (Leica). For the 2×8 mm bioprinted disks we proceeded directly to follow immunofluorescence staining without sectioning. For the immunofluorescent labeling of the constructs, the samples were permeabilized with 0.1% (v/v) Tween-20 for 1 h at 4°C. After permeabilization, the constructs were blocked with a blocking buffer (0.1% (v/v) Triton X-100 and 1% (w/v) BSA in PBS) for two hours. After blocking, the constructs were incubated overnight at 4°C with primary antibodies goat polyclonal anti-Pdx1 (1:300 dilution, R&D systems) and rabbit monoclonal anti-MafA [D2Z6N] (1:800, Cell Signaling Technology) diluted in blocking buffer. After incubation with the primary antibodies the samples were washed for 15 min three times with blocking buffer and then incubated overnight at 4°C in the dark with secondary antibodies donkey anti-goat AlexaFluor 488 (1:300 dilution, Thermo Fisher Scientific), donkey anti-rabbit AlexaFluor 555 (1:300 dilution, Thermo Fisher Scientific), and 1 μg/mL Hoechst 33342 (Thermo Fisher Scientific) diluted in blocking buffer. Samples were then washed for 15 min three times with PBS and examined on a Leica Thunder microscope.

### Pdx1 and MafA expression quantification from immunofluorescence microscopy images

To quantify MafA and Pdx1 levels from immunofluorescence stainings, raw images were exported as 16-bit grayscale. For 3D cultures, maximum intensity projections were generated using Leica Application Suite X (v5.3.1). Background correction was performed in ImageJ, followed by automated quantification of nuclear counts and background-subtracted mean intensities for MafA and Pdx1 using a custom macro. Nuclear segmentation was performed on Hoechst-stained images via manual thresholding and watershed separation, with nuclei enumerated using particle analysis. Three background regions of interest (ROIs) were manually selected per image and channel, and mean fluorescence intensities were measured across the full image and background ROIs. Corrected fluorescence intensities were calculated by subtracting background values, and per-nucleus expression levels were derived by normalizing corrected intensities to nuclear counts. As a complementary strategy, protein expression was assessed by calculating the fraction of nuclei classified as MafA⁺ or Pdx1⁺ above a defined intensity threshold. Each marker channel was processed using ImageJ’s default thresholding algorithm, with the cutoff set at the highest value that eliminated speckling, then binarized and separated by watershed. Particle analysis with size (>15 pixels) and circularity (≥0.40) constraints excluded artifacts, and marker-positive counts were expressed as a percentage of total nuclei. To assess co-localization between MafA and Pdx1, CellProfiler (v4.2.6, Broad Institute) was used to calculate Pearson’s correlation and Manders’ overlap coefficients with the MeasureColocalization module. To visualize the distribution of fluorescence along the height of the constructs, intensity side profiles were generated using ImageJ’s plot profile function. Raw data arrays were imported into a Python (v3.11), binned in 200 µm intervals to eliminate local irregularities, and min–max normalized per channel. Normalized data and binned means were exported to Excel, and profiles were visualized as line plots per channel. As a measure of spatial selectivity, the illuminated target region was overlayed on the side profile, and the % of the full AUC within this domain was calculated using the previously python script.

### Live/Dead assay for cell viability analysis

Microencapsulated and volumetrically bioprinted HeLa mEos3.2 and volumetrically printed transfected HeLa cells were stained with Calcein AM and ethidium homodimer-1 to determine the cell viability. The constructs were incubated in a PBS solution containing 4 µM ethidium homodimer-1 and 2 µM Calcein AM for 30 mins at ambient temperature in the dark. Subsequently, the constructs were washed three times for 10 min with PBS and then imaged on a Leica Thunder microscope.

### mRNA expression quantification of PDX1 and MAFA via RT-qPCR

After illuminating HeLa cells with NIR as described previously, RNA was isolated using the RNeasy kit (Qiagen, 74104), according to the manufacturer’s protocol. For cDNA synthesis, 500 ng of RNA was combined with the components of the cDNA synthesis kit iScript (Biorad) and thermocycling was carried out following the manufacturer’s instructions. RT-qPCR was performed using an iQ SYBR Green Kit (Biorad) following the manufacturer instructions. Briefly, a primer mix was prepared by dissolving 1 µL of each gene-specific forward and reverse primer (100 µM stock) in 28 µL of nuclease-free water. Primer sequences as follows: *PDX1* forward: AGCTGCCCTTTCCTTGGATG, *PDX1* reverse: TATAGGCTGTCCGGGTTCGT, *MAFA* forward: CAGCACCATCTGAACCCTGA, *MAFA* reverse: CCAGCTCCCATATCGTCTGC, *GAPDH* forward: CAACGGATTTGGTCGTATTGGG, *GAPDH* reverse: TGCCATGGGTGGAATCATATTGG. The qPCR was run on the CFX96™ Real-Time PCR System with an initial 95 °C denaturation (3 min), followed by 40 cycles of 95 °C for 10 s and 60 °C for 45 s. Data were analysed using the comparative Ct (ΔΔCt) method, normalizing target gene expression to *GAPDH* as housekeeping gene.

### Statistical analysis

Results were reported as mean ± standard deviation (S.D.). Statistical analysis was performed using GraphPad Prism 8 (GraphPad Software, USA). Comparisons between multiple experimental groups were assessed via one or two-way ANOVAs, followed by post hoc Bonferroni correction to test differences between groups. Student’s t-test was performed between 2 experimental groups for statistical analysis. An unpaired t-test was used for parametric comparisons of data sets. Non-parametric tests were used when normality could not be assumed. Pdx1 and MafA signal correlation was assessed by a Meanders’ correlation coefficient. RT-qPCR data were analyzed using the comparative Ct (ΔΔCt) method, normalizing target gene expression to *GAPDH* as housekeeping gene, and a paired t-test was used to assess significance on paired samples coming from the same transfections rounds (illuminated vs dark control). Differences were considered significant when *p* < 0.05.

## ACKNOWLEDGMENTS

D.R., and P.C. contributed equally and share first authorship. G.G. and P.N.B. contributed equally. This project received funding from the European Research Council (ERC) under the European Union’s Horizon 2020 research and innovation programme (grant agreement No. 949806, VOLUME-BIO) and under the European Union’s Horizon Europe research and innovation programme (grant agreement No. 101231578, SMART-AGENT), and from the European’s Union’s Horizon 2020 research and innovation programme under grant agreement No 964497 (ENLIGHT). Views and opinions expressed are however those of the author(s) only and do not necessarily reflect those of the European Union or the European Research Council. Neither the European Union nor the granting authority can be held responsible for them. W.N and L.C.K. were supported by the EWUU Alliance, via the Centre for Living Technologies. R.L. and J.M. acknowledge the funding from the Gravitation Program “Materials Driven Regeneration” (024.003.013) and from the Summit program “DRIVE-RM” (SUMMIT.1.027), funded by the Netherlands Organization for Scientific Research (NWO). R.L. also acknowledges financial support from the Dutch Research Council (Vidi, 20387).

## DATA AVAILABILITY STATEMENT

The data that support the findings of this study are available from the corresponding author upon reasonable request.

## Supplementary Information

## SUPPORTING INFORMATION

### Supplementary methods

#### Optical density measurements for GelMA and AlgMA

GelMA (5% w/v) and AlgMA (3% w/v) solutions were prepared by dissolving the pre-polymers in PBS, and supplemented with 0.1% w/v LAP, and either mixed with HeLa cells (2 million/mL) or left cell-free. Cell-laden and cell-free suspensions were placed in a 96-well multiwell plates (250 uL/sample), and photocrosslinked (405nm, 3.1 mW/cm2, 5 minutes). Pre-crosslinked, cell-free solutions were also added, together with a PBS-only control. All groups were included with 3-4 replicates. Optical density at 405nm was measured with a spectrophotometer (CLARIOstar Plus, BMG Labtec), and absorbance values were recorded.

##### Alginate photoactivation assays

Alginate methacrylate was dissolved in PBS to 3% (w/v) and mixed with HeLa mEos3.2 cells at 2 × 10⁶ cells/mL. The resulting bioink was cast under a 405 nm lamp for 2 minutes (delivering a light dose of 372 mJ/cm^2^) and transferred to a cuvette. A dose test was then performed on a volumetric printer (Tomolite v2, Readily3D, Switzerland) using its Apparite software, by projecting a matrix of circular spots each illuminated at a different light dose (1000, 2500, 5000, 7500, 14000, and 20000 mJ/cm²) obtained with variable light intensity (20, 30, 40, and 50 mW/cm²) (n = 3).

#### Characterization of the support bath for Embedded extrusion Volumetric Printing

To perform as a suitable bath for embedded extrusion printing, the low molecular weight GelMA needs to allow for the movement of the immersed needle, while offering structural support to the extruded filament and while preserving its stress-recovery capacity. To characterize and confirm this behaviour, a shear strain sweep test was initially performed, and results showed a cross point of *G*′ and G″ at 440% strain, resulting in a solid-liquid transition (**Figure 3B**, in the main body of the manuscript). Secondly, a shear rate ramp was performed and a shear thinning behaviour was observed as the shear stress increased in a nonlinear fashion and, under the same conditions, viscosity decreased (**Figure 3C**). Finally, stress/recovery tests were performed by alternating low (1%) and high (500%) strain at 1 Hz frequency. Results showed how, when the higher strain was applied, higher G″ values were measured compared to the *G*′ ones, (**Figure 3D**), proving how the fluid bulk support bath transitioned from a solid to a liquid-like state when a strain was applied, representing the needle translation through the bath during the extrusion phase. Moreover, higher *G*′ values were registered compared to *G*″ after the strain removal, showing an elastic recovery, thus proving the ability of support for the extruded filament. Following the printability optimization, filaments of width ranging from 331 ± 23 µm to 134 ± 22 µm were extruded with a methylcellulose-based ink, previously designed and characterized.^[6]^ With the range of width values achieved, a filament smaller or bigger than the diameter of the needle (27G) utilized in this study could be achieved (**Figure 3E**).

#### Cell viability of primary stem cells under photoconversion light dose regimes

Mesenchymal stromal cells (MSCs) were isolated from equine bone marrow as previously described, [SR1] and cultured in αMEM (Gibco) supplemented with 0.2 mM L-ascorbic acid 2-phosphate (Sigma), 10% FCS (Lonza), 100 U/mL penicillin with 100 µg/mL streptomycin (Life Technologies) and 1 ng/mL bFGF and used at passage 5. All animal tissue and cells used in this study were obtained from deceased equine donors, donated to science by their owner, and according to the guidelines of the Institutional Animal Ethical Committee. 2×10^6^ MSCs were volumetrically printed in 5% GelMA into cylindrical samples (2mm height, 6 mm diameter) at a light dose of 180 mJ/cm^2^. Samples were kept in culture overnight to wash out any excess photoinitiator and subsequently exposed to additional light irradiation to mimic the highest photoactivation regimes used on the HeLa mEos3.2 experiments (11.25 and 11.73 J/cm^2^, for static and tomographic 3D photoactivation, respectively). Irradiated samples, and non-irradiated controls (n = 3) were cultured for 3 days. Live/Dead assays were performed according to manufacturer’s protocols (ThermoFisher) at days 1 and 3 to assess cell viability post-irradiation. Samples were imaged using a Thunder microscope (Leica) and percentage of live cells calculated using FIJI.

#### Assessment of the temporal expression of PDX1 and MAFA after NIR illumination

HeLa cells were plated in 96-well plates at a cell density of 10000 cells/well. The day after plating, the medium was refreshed, and HeLa cells were transfected with both plasmid constructs as previously described. 24 h post transfection, 2D transfected HeLa cells were exposed to pulsed light regimes of 30 s ON/180 s OFF using 740 nm LED (Roithner, LED740-01AU), 25 mW/cm^2^, connected to an Arduino Mega (microcontroller board, Arduino) via a LED DRIVER (Infineon ILD6150), in phenol red-free cell culture media supplemented with 25 µM biliverdin (30891, Sigma-Aldrich) overnight. Cells were analyzed at different timepoints after induction of light exposure via immunofluorescence staining as described below. Following optogenetic activation 2D-cultured cells were fixed in 4% formaldehyde solution for 15 min at room temperature and permeabilized with 0.1% (v/v) Triton X-100 in phosphate buffered saline for 10 min. After permeabilization, cells were blocked with a blocking buffer (1% (w/v) BSA solution in PBS) for 1 h at room temperature. The cells were incubated overnight at 4°C with primary antibodies goat polyclonal anti-Pdx1 (1:300 dilution, R&D systems) and rabbit monoclonal antiMafA [D2Z6N] (1:800, Cell Signaling Technology) diluted in blocking buffer. After primary antibody incubation the cells were washed three times with blocking buffer and then incubated with secondary antibodies donkey anti-goat AlexaFluor 488 (1:300 dilution, Thermo Fisher Scientific), donkey anti-rabbit AlexaFluor 555 (1:300 dilution, Thermo Fisher Scientific), and 1 μg/mL Hoechst 33342 (Thermo Fisher Scientific) diluted in blocking buffer for 1 h at room temperature in the dark. Samples were then washed three times in PBS and analyzed on a Leica Thunder inverted microscope. Signal quantification was performed in Fiji (ImageJ) by threshold-based segmentation of Hoechst-stained nuclei followed by watershed separation to determine nuclei counts per image.

#### Viability of HeLa cells after transfection and encapsulation

HeLa cells were plated in 6-well plates at a cell density of 800000 cells/well. The day after plating, the medium was refreshed, and HeLa cells were transfected with both plasmid constructs as previously described. 24h post transfection, cells were encapsulated into 6 x 2 mm GelMa discs at a cell density of 2e6 cells/mL. Cell viability was assessed 24h and 72h post encapsulation by staining the constructs with 2 µM Calcein AM and 4 µM ethidium homodimer-1 (EH) for 30 min at room temperature in the dark, and imaged on a Leica Thunder microscope. Live-dead cell quantification was performed in Fiji (ImageJ). Images were treated with a sharpen filter to facilitate by threshold-based segmentation applied independently to the Calcein AM and EH channels, followed by watershed separation to identify and count individual cells. The numbers of live (Calcein AM-positive) and dead (EH-positive) cells were determined for each image, and cell viability was calculated as the percentage of live cells relative to the total number of cells (live + dead).

### Supplementary results

#### Photostimulation of HeLa-mEos3.2 embedded in AlgMA

As a proof-of-concept demonstration of applying our approach to different classes of biomaterials, we tested Alginate Methacryloyl (AlgMA) as a polysaccharide-based bioresin.(Supplementary Figures S3, S4, and S5). Also in AlgMA, we observed how the dose needed for printing (approximately 372 mJ/cm^2^) is considerably lower (approximately 3-fold) than the minimum dose required to activate the mEos3.2 protein, and that the efficiency of photoconversion increases with the applied dose. For the AlgMA-based bioresin, a spatial activation accuracy of 84 ± 3%, with an efficiency of 72 ± 4% of cells converted were measured. As additional characterization, we assessed also the optical density of the AlgMA-bioresin at 405nm, which increased approximately 3-fold over cell-free hydrogels when cells were embedded. This is in line with the increased opacity of light-scattering, cell-laden structures. Although the overall optical density of crosslinked AlgMA 3% was found to be significantly lower (p<0.05, t-test, n=3) than what we measured for our GelMA 5%-based formulations, a direct comparison should be considered with caution, as the two materials are vastly different in composition, polymer content, molecular weight and degree of functionalization. Overall, our data underlines that photoactivation is also possible with this alginate-based material.

### Supplementary figures

**Supplementary Figure S1:**
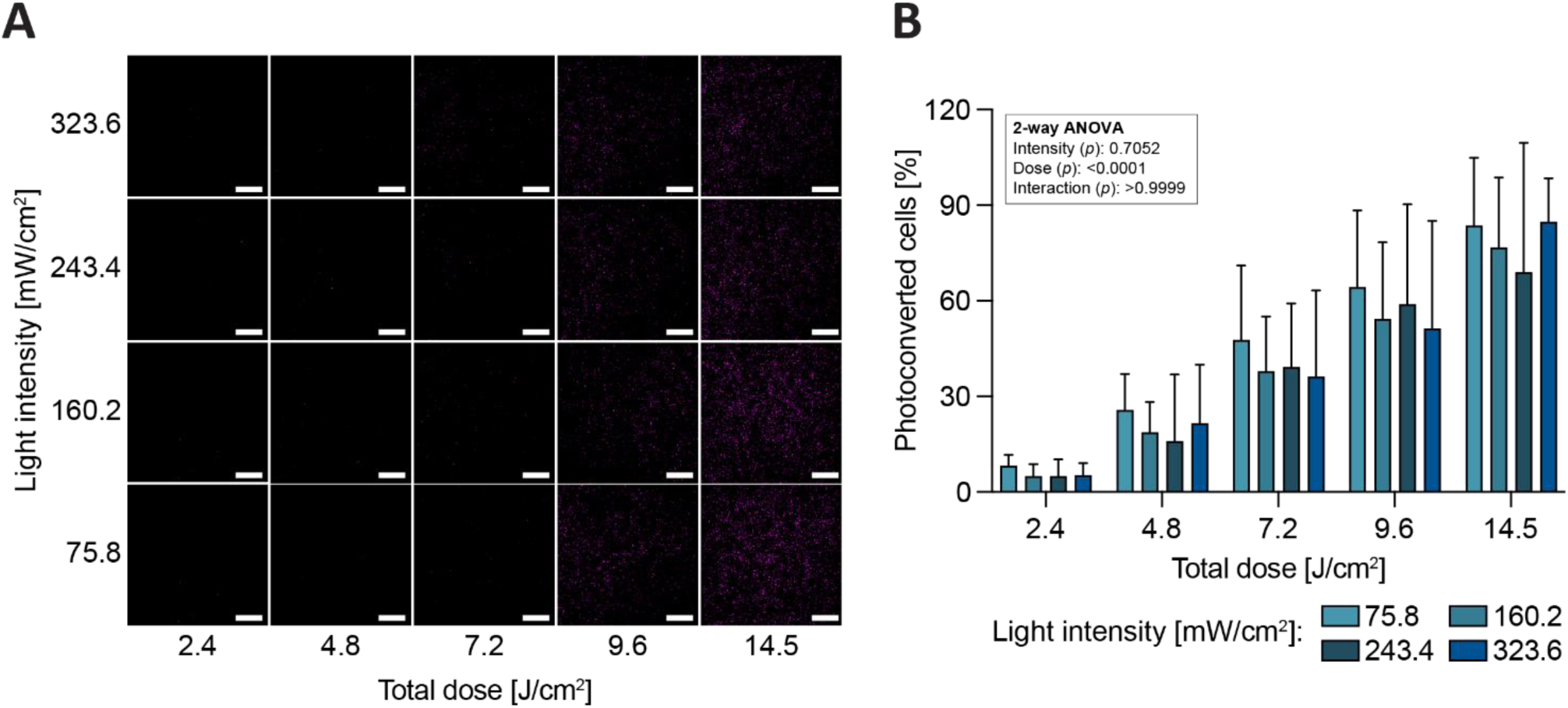
A) Representative fluorescence images of photoconverted HeLa mEos3.2 (magenta) exposed to the complete grid of screened dose and light intensity conditions for the characterization of the dose dependency behaviour of mEos3.2 photoconversion in 2D. Scale bar: 500 µm. B) Plotted percentage of photoconverted cells in each of the tested conditions (n = 3) and results of the two-way ANOVA showing that dose is the only significant factor affecting photoconversion efficiency.

**Supplementary Figure S2:**
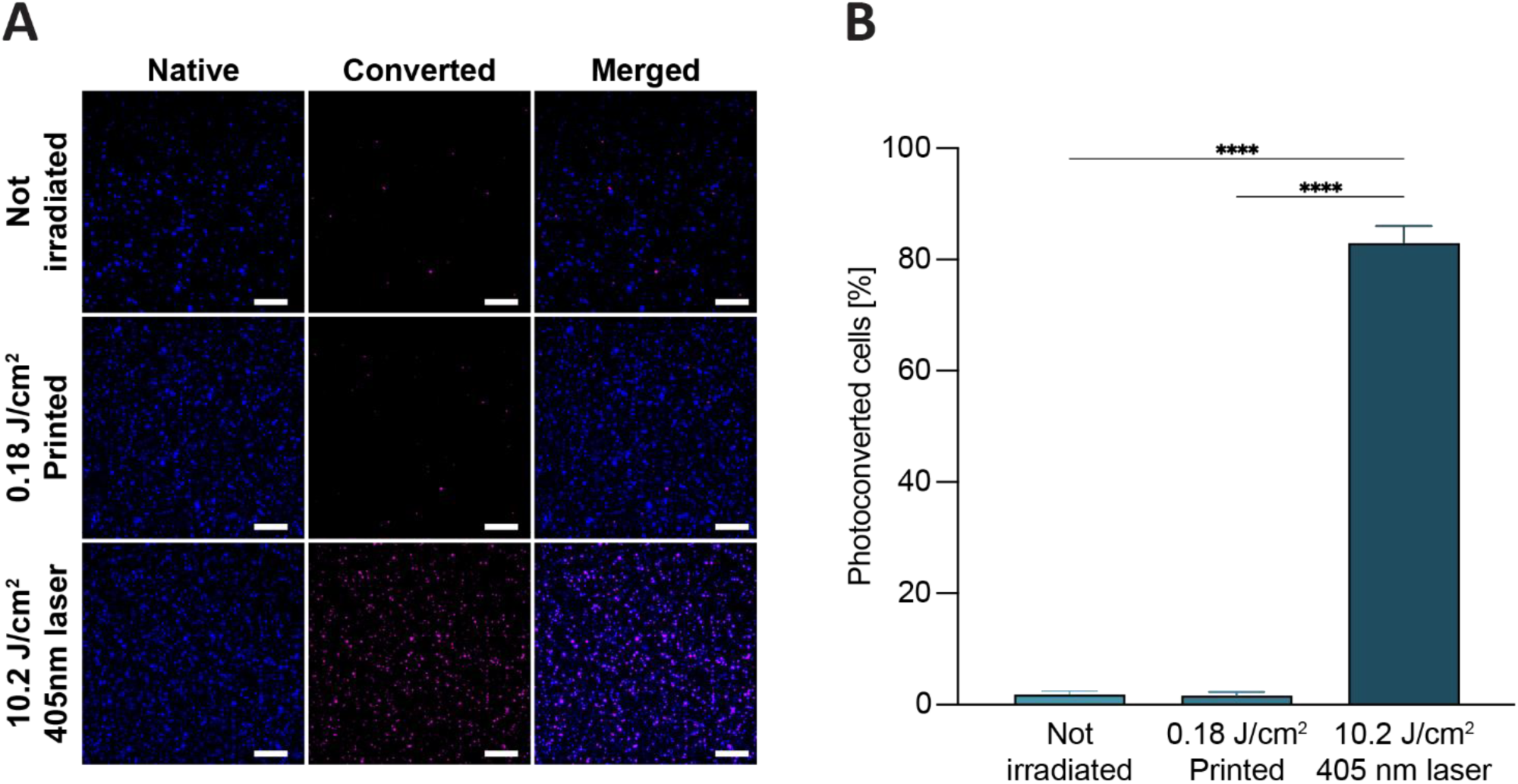
A) Fluorescence images and B) percentage of native (blue) and photoconverted (magenta) HeLa mEos3.2 cells pre-treated with doxycycline and embedded in 5% GelMA i) before crosslinking (not irradiated), ii) crosslinked in the volumetric bioprinter at a light dose of 180 mJ/cm2, and iii) fully converted at a dose of 10.2 J/cm2 using the 405 nm laser of a Thunder microscope (n = 5). Scale bar: 200 µm. Data represent mean ± SD. Significance was determined by one-way ANOVA: **** = p< 0.0001.

**Supplementary Figure S3:**
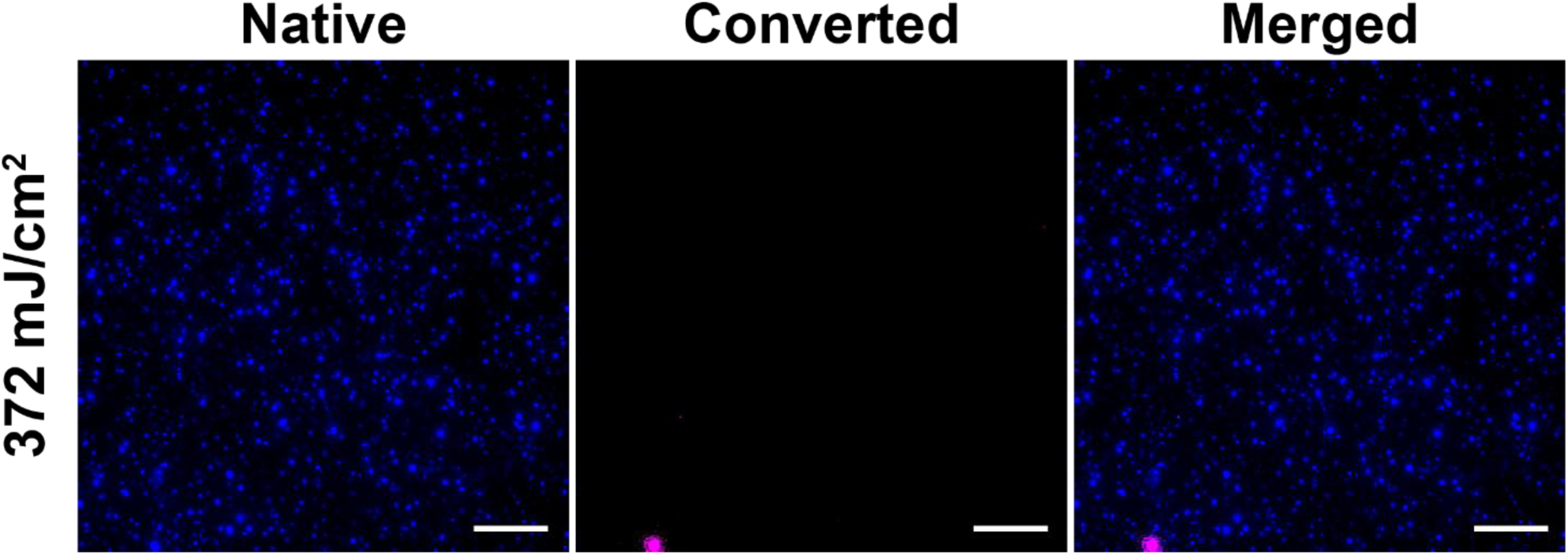
Fluorescence images of native (blue) and photoconverted (magenta) HeLa mEos3.2 cells pre-treated with doxycycline and embedded in 3% AlgMA via photocrosslinking at a dose of 372 mW/cm^2^, demonstrating that the light dose required to crosslink the cell-laden material does not lead to photoconversion. Scale bar: 200 µm.

**Supplementary Figure S4:**
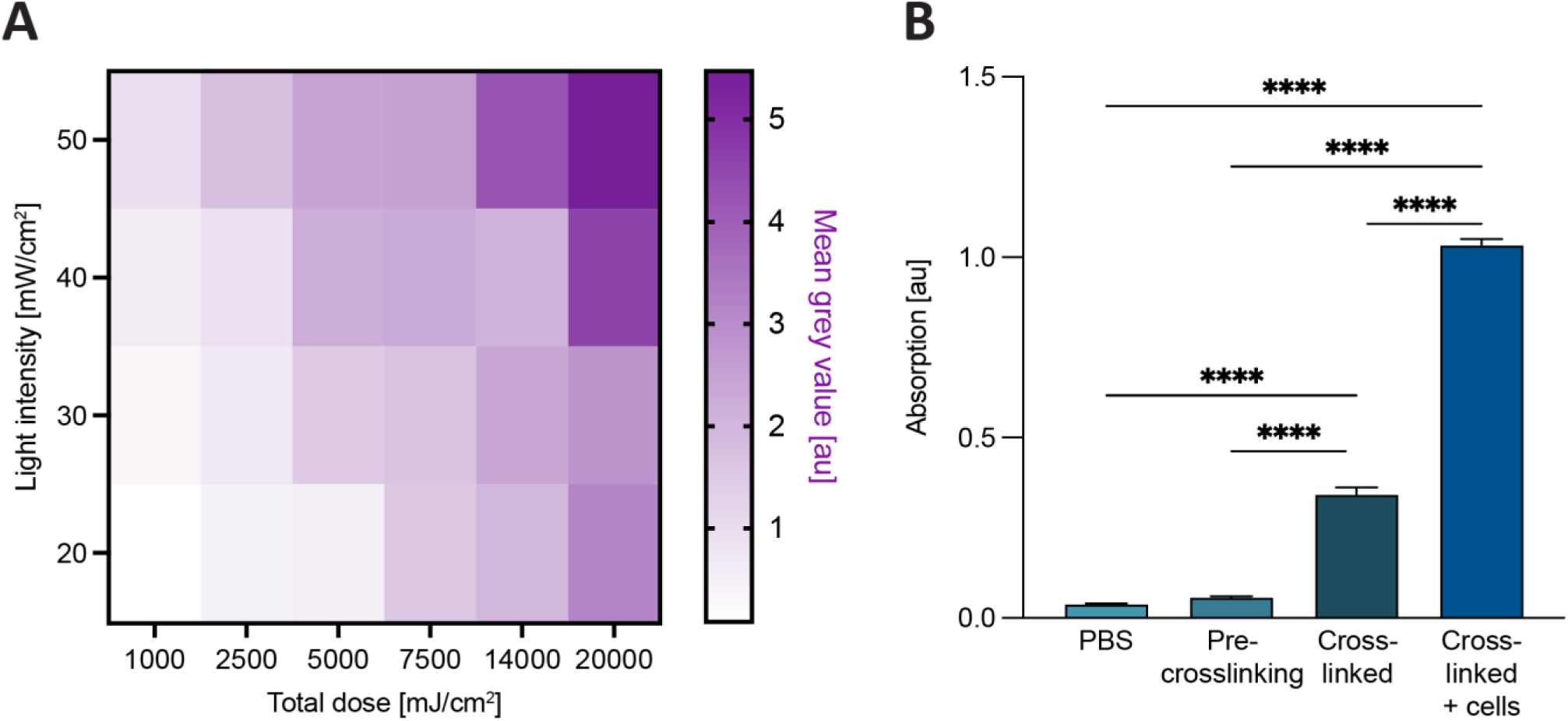
A) Screening of volumetric printing set-up light intensity and total light dose required for HeLa-mEos3.2 photoconversion in 3% AlgMA samples (n = 3). B) Optical density at 405nm for AlgMA solutions, cell-free and cell-laden crosslinked hydrogels. Data is represented as mean ± SD. Significance was determined by one-way ANOVA: **** = p< 0.0001.

**Supplementary Figure S5:**
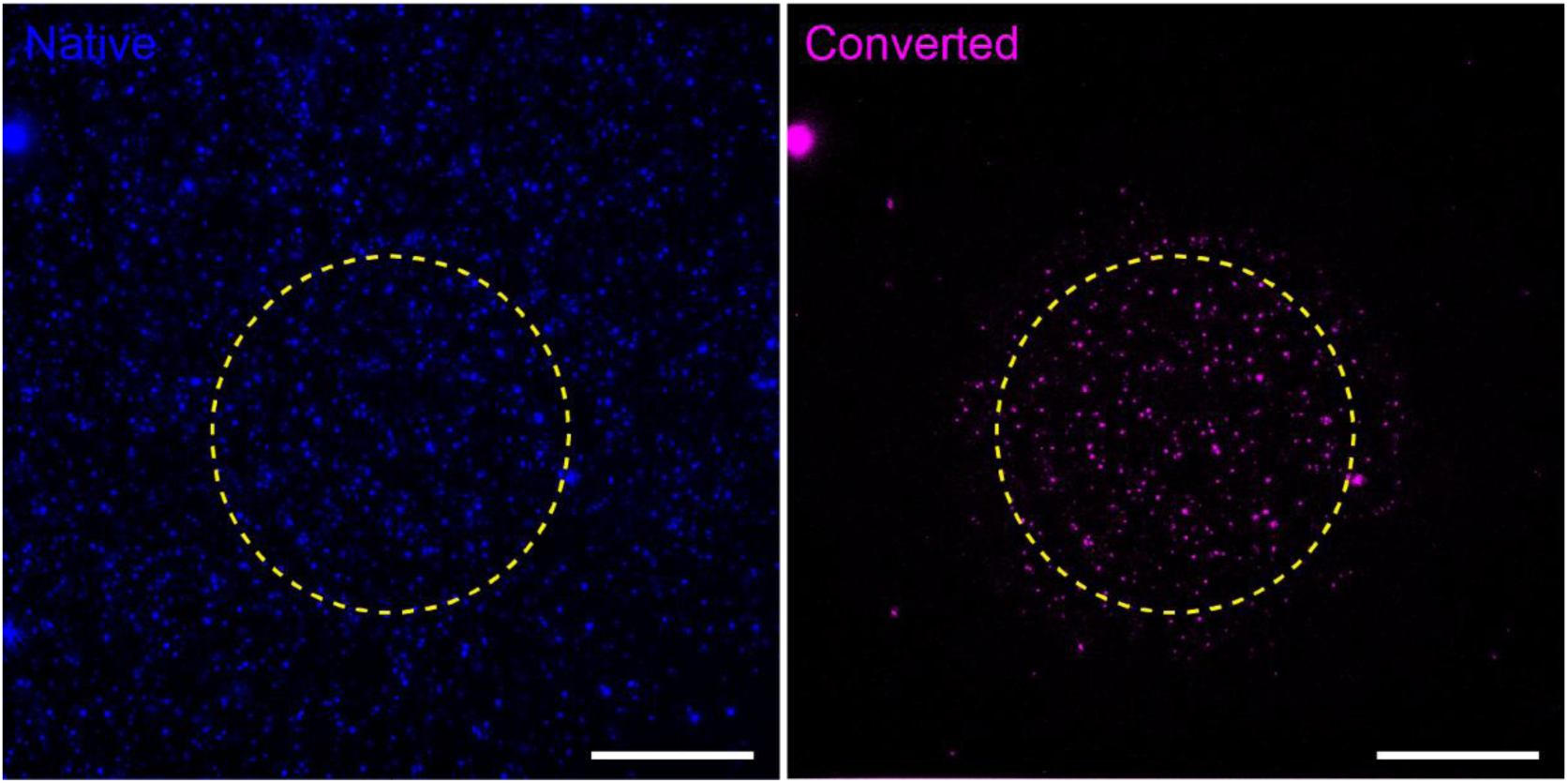
A) Photoconversion accuracy in AlgMA. Scale bar: 500 µm.

**Supplementary Figure S6:**
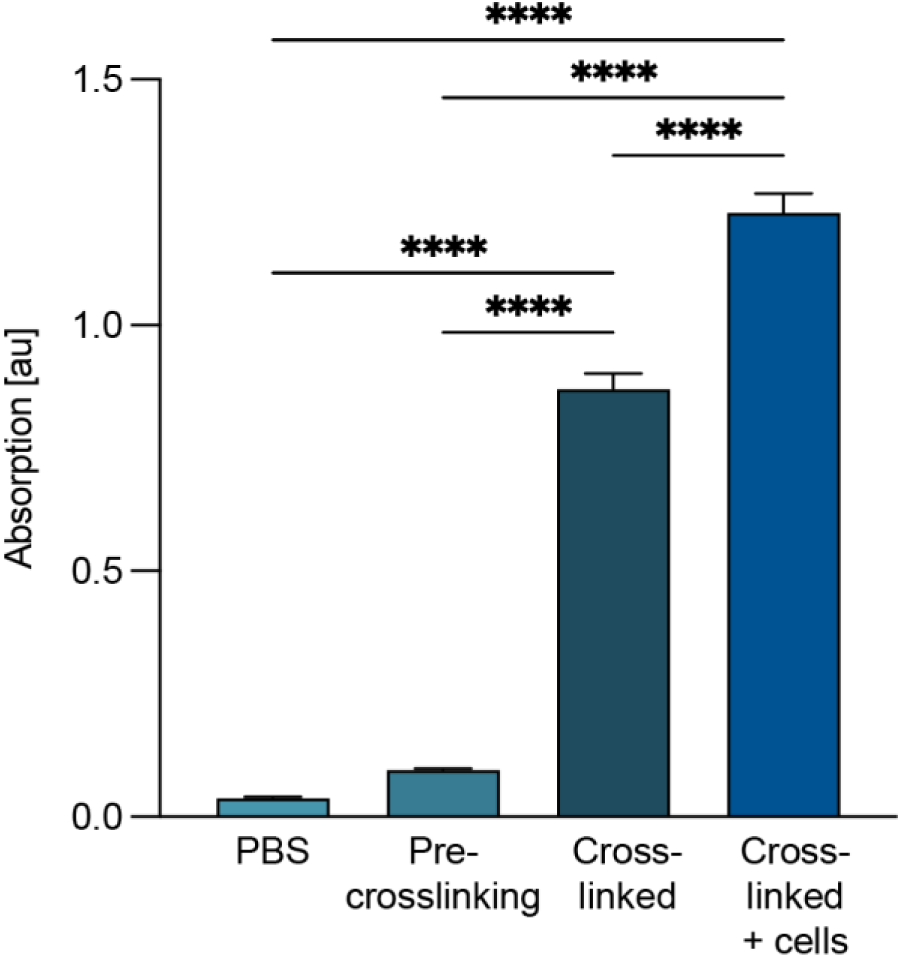
Optical density at 405m for GelMA solutions, cell-free and cell-laden crosslinked hydrogels. Data is represented as mean ± SD. Significance was determined by one-way ANOVA: **** = p< 0.0001.

**Supplementary Figure S7:**
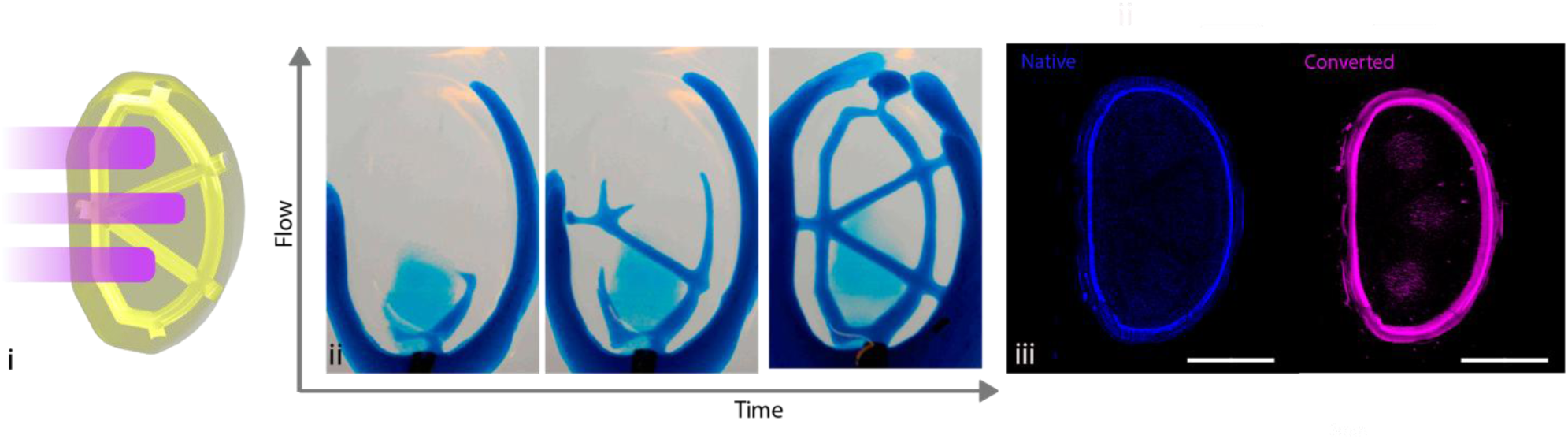
3D photoconversion of HeLa mEos3.2 in a bioprinted lymph node-shaped model (volume = 190 mm^3^). i) STL model showing the printed, perfusable lymph structure (beige/yellow) and the desired photoactivated regions (magenta). ii) Manual perfusion of the volumetrically printed channels with an alcian blue dye. iii) Fluorescence imaging tilescan of the whole construct showing the photoconverted regions. Scale bar: 3 mm.

**Supplementary Figure S8:**
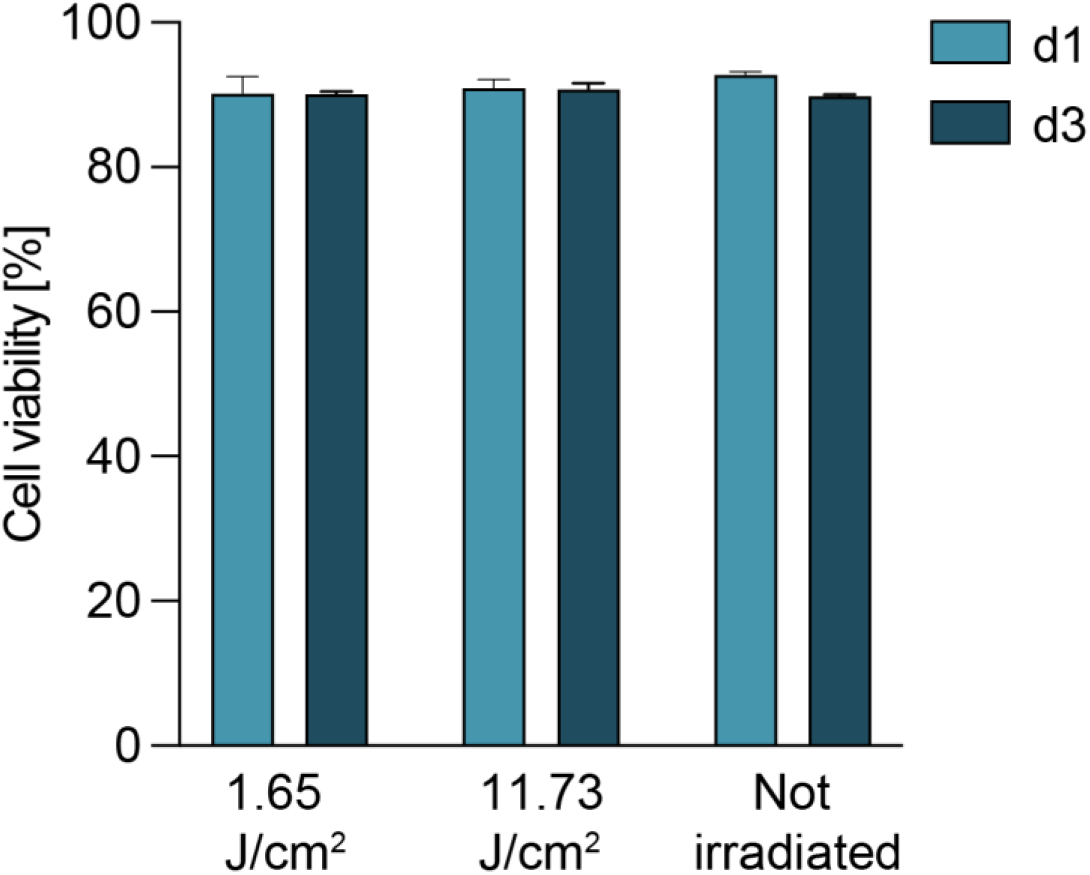
Cell viability at day 1 and 3 of primary mesenchymal stromal cells from equine bone marrow printed within 5% GelMA (180mJ/cm2 dose) and subsequently irradiated at a low dose, and a dose just above the highest photoactivation regime used on the HeLa mEos3.2 experiments (11.73 J/cm^2^) using a volumetric bioprinter (n = 3). Data represent mean ± SD. No significant differences were determined by two-way ANOVA.

**Supplementary Figure S9:**
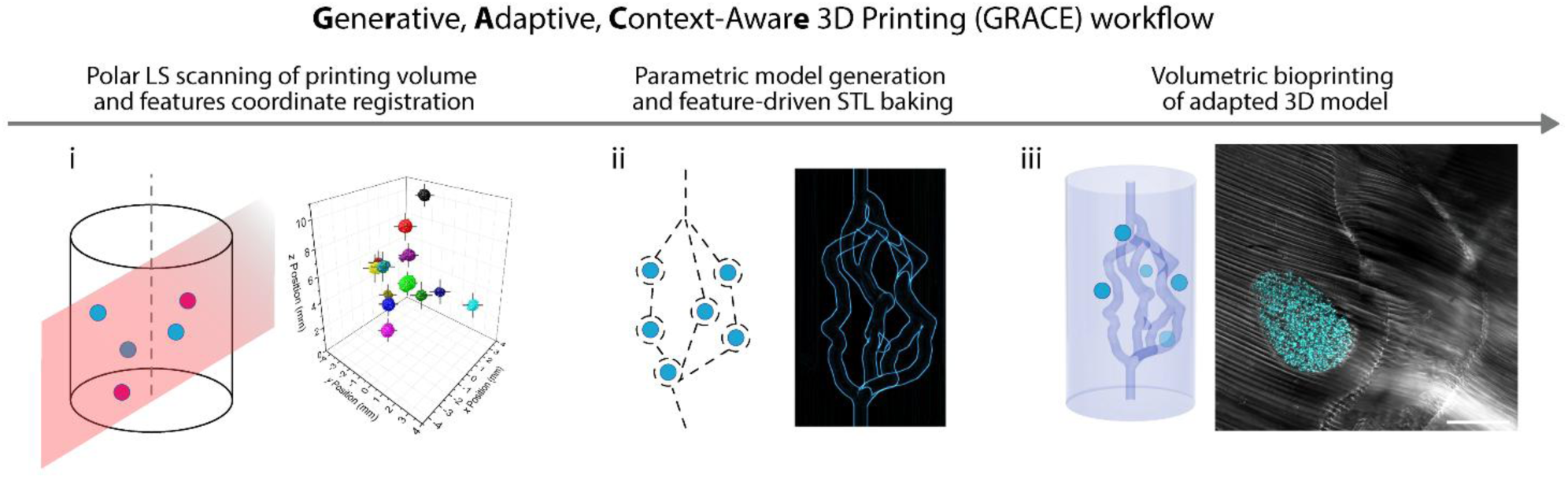
GRACE workflow schematics. Briefly, a light sheet rapidly scans and maps the printing volume in 3D to extract positional, morphometric, and spectral (i.e., fluorescence) information from the contents of the printing vial. This serves as input for multi-parametric modelling algorithms to generate precisely a targeted geometry, which is finally converted into the STL model to be volumetrically printed. Scale bar for panel iii: 500 µm.

**Supplementary Figure S10:**
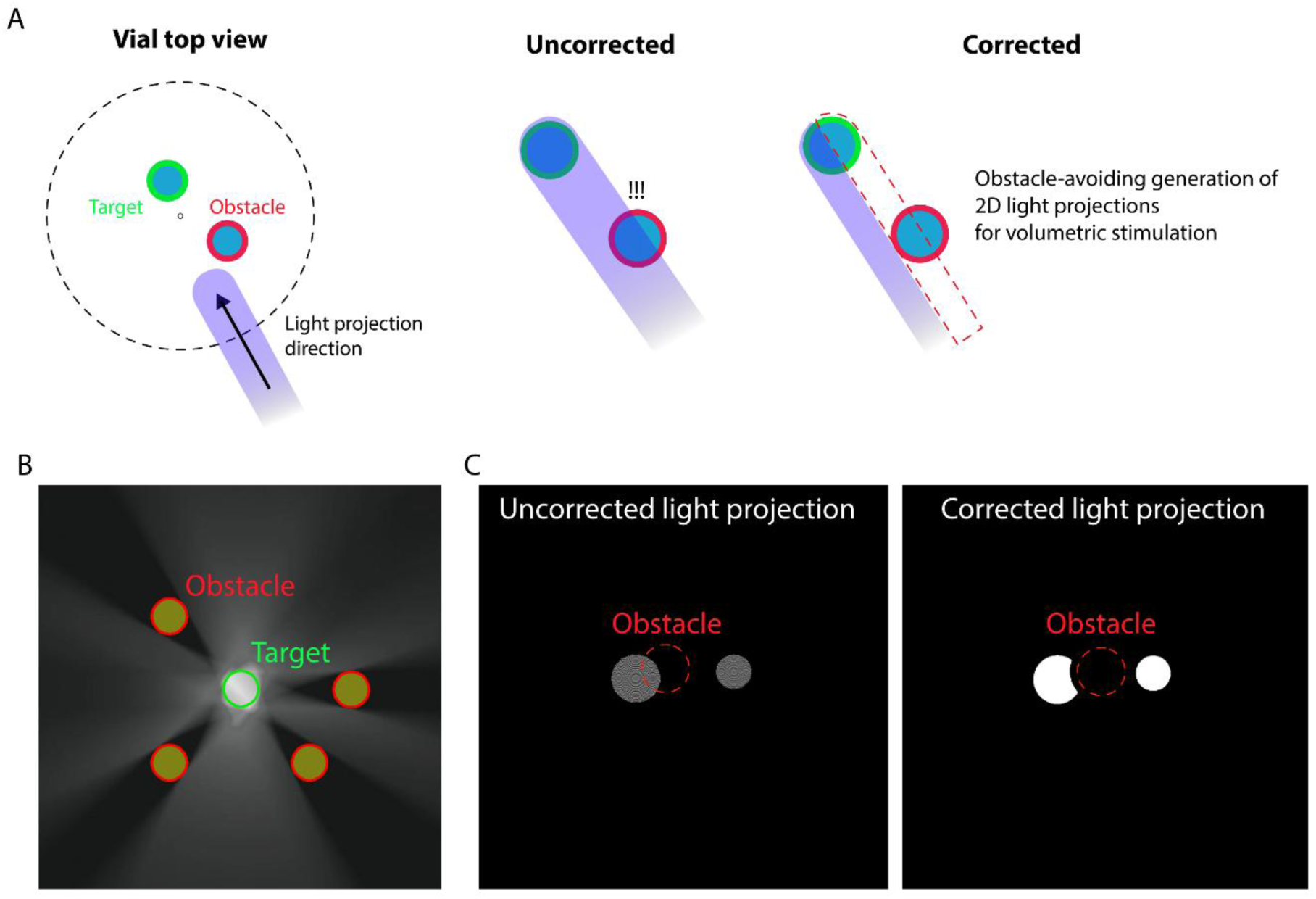
GRACE obstacle correction schematic explanation for the selective photoconversion of HeLa-mESO3.2.

**Supplementary Figure S11:**
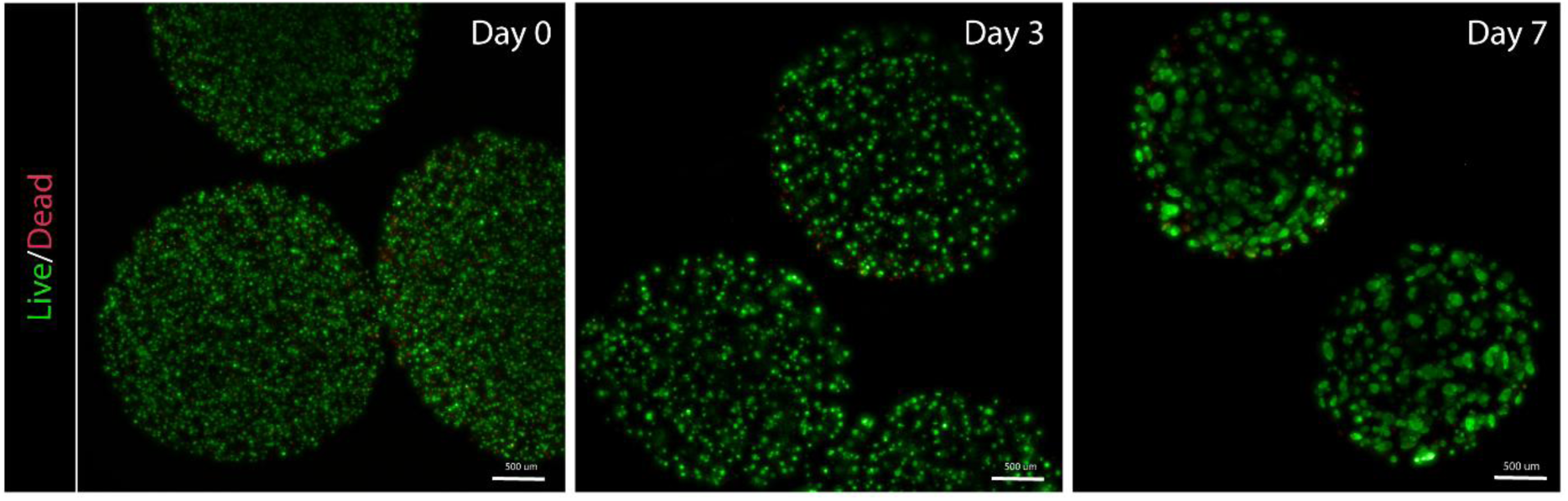
Live/Dead essay on encapsulated HeLa cells over a period of 7 days.

**Supplementary Figure S12:**
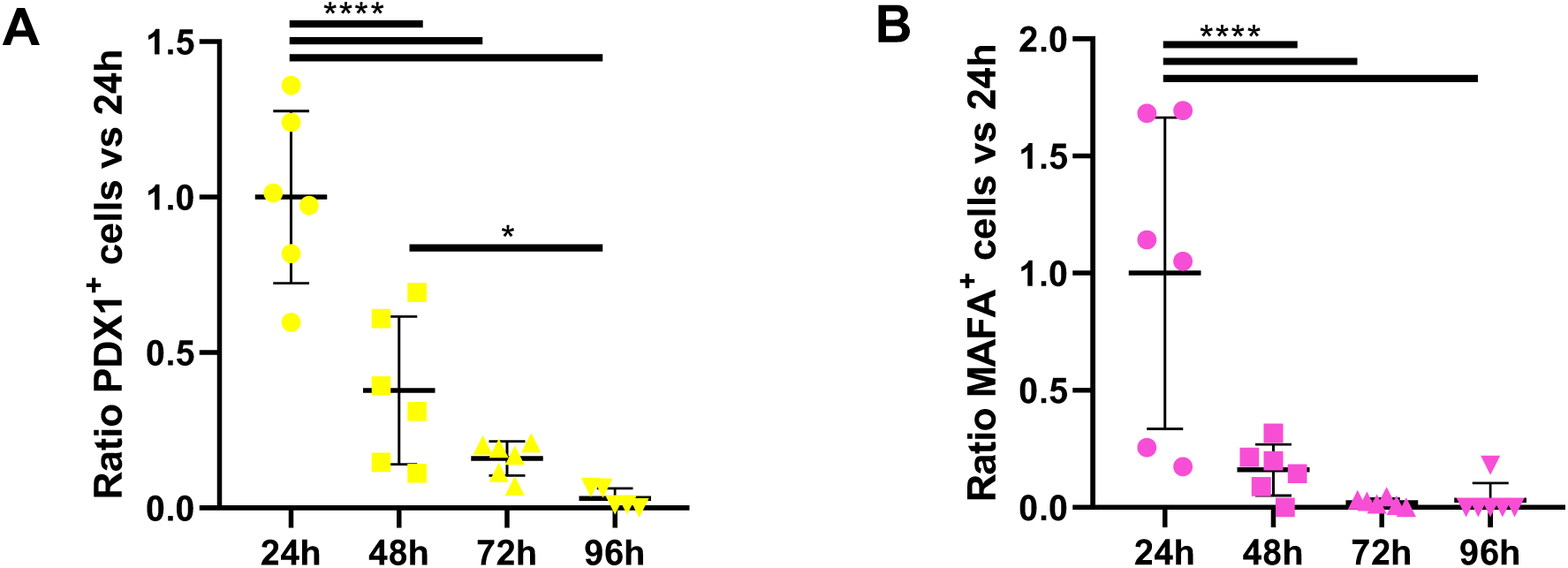
Recovery of baseline protein expression over time after photostimulation of the NIR-sensitive circuit, for A) PDX1 and B) MAFA, normalized to the average of the expression level found 24h after induction of photostimulation (n=6, * p = 0.021, **** p <0.0001, one-way ANOVA).

**Supplementary Figure S13:**
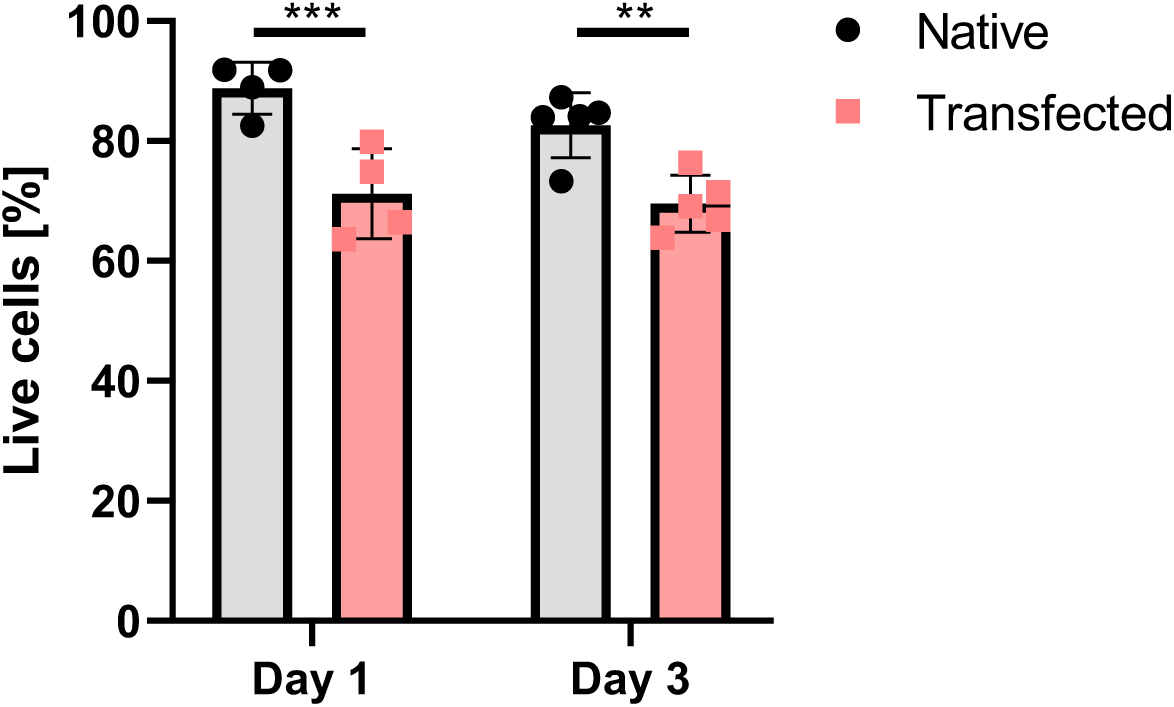
Viability of HeLa cells in their native state or after transfection with the NIR-sensitive circuit, after embedding in GelMA. Transfected cells display lower viability compared to non-transfected, albeit the overall percentage of living cells is ⁓70% (n=4-5, ** p = 0.042, *** p = 0.0008, two-way ANOVA).

**Supplementary Figure S14.**
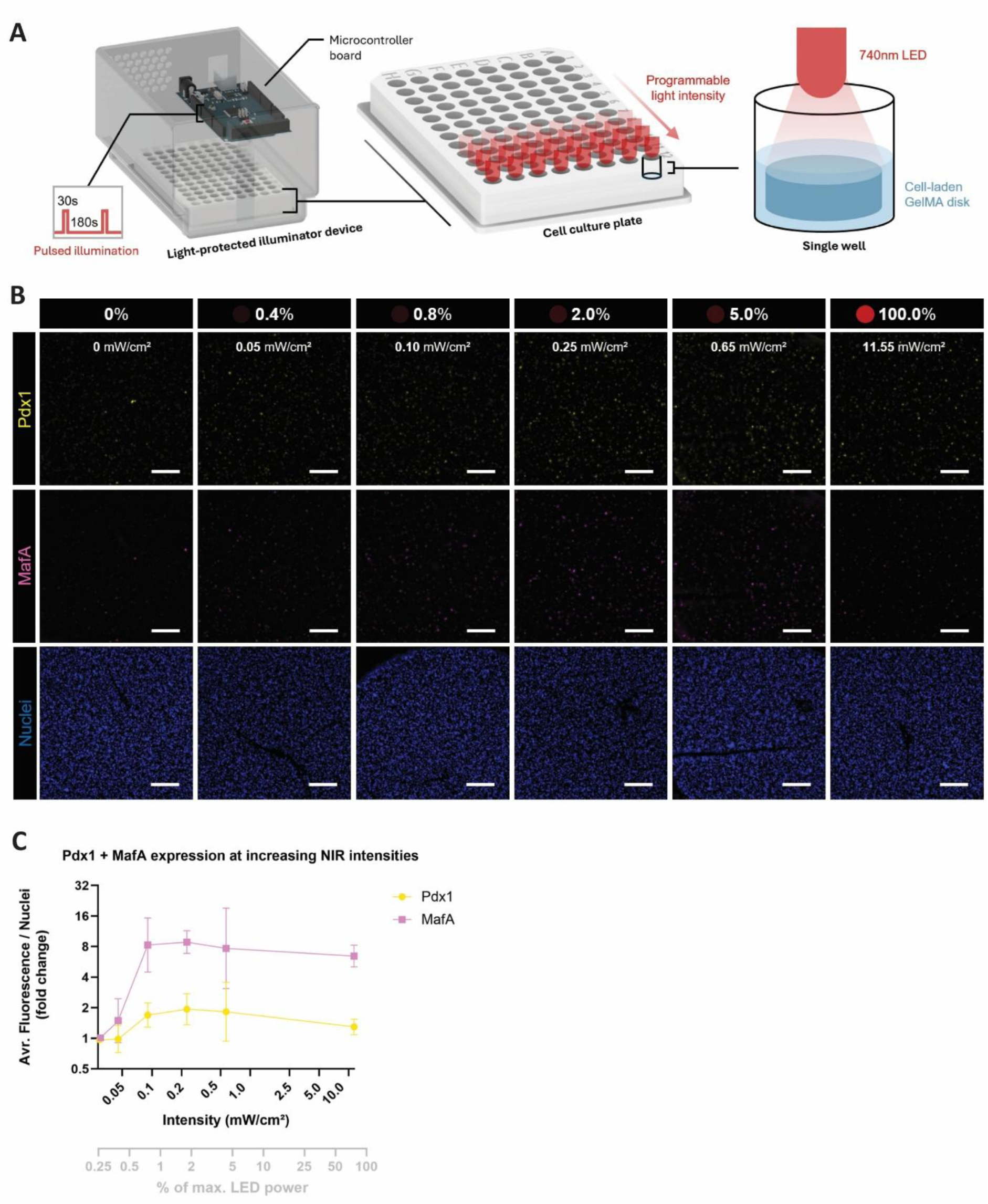
NIR optogenetic system displays intensity-dependent activation in 3D cultures. (A) A custom illuminator device was built to illuminate the wells of a 96-well plate with programmable light intensity. A microcontroller controlled a custom illumination script consisting of 30 s ON and 180 s OFF cycles, with the average intensity independently adjustable for each row of wells. To shield samples from ambient light, the plate was enclosed in a black 3D-printed box. In each well, a 4 × 1 mm GelMA disks was placed, submerged in medium, and illuminated according to the script for 24 h. (B) 5× IF stainings of Pdx1/MafA expression in GelMA disks after illumination at increasing light intensities. Top values represent % of total LED power, whereas bottom values are absolute intensities. Nuclei are visualized with HOECHST. S.B. = 500 μm. (C) Quantification of Pdx1/MafA expression based on average (avr) fluorescence by cell nuclei count in GelMA disks after illumination at increasing light intensities. Y-axis displays average image fluorescence per number of nuclei, normalized to the unilluminated control (N = 1, n = 3). Data represent mean ± SD.

**Supplementary Figure S15.**
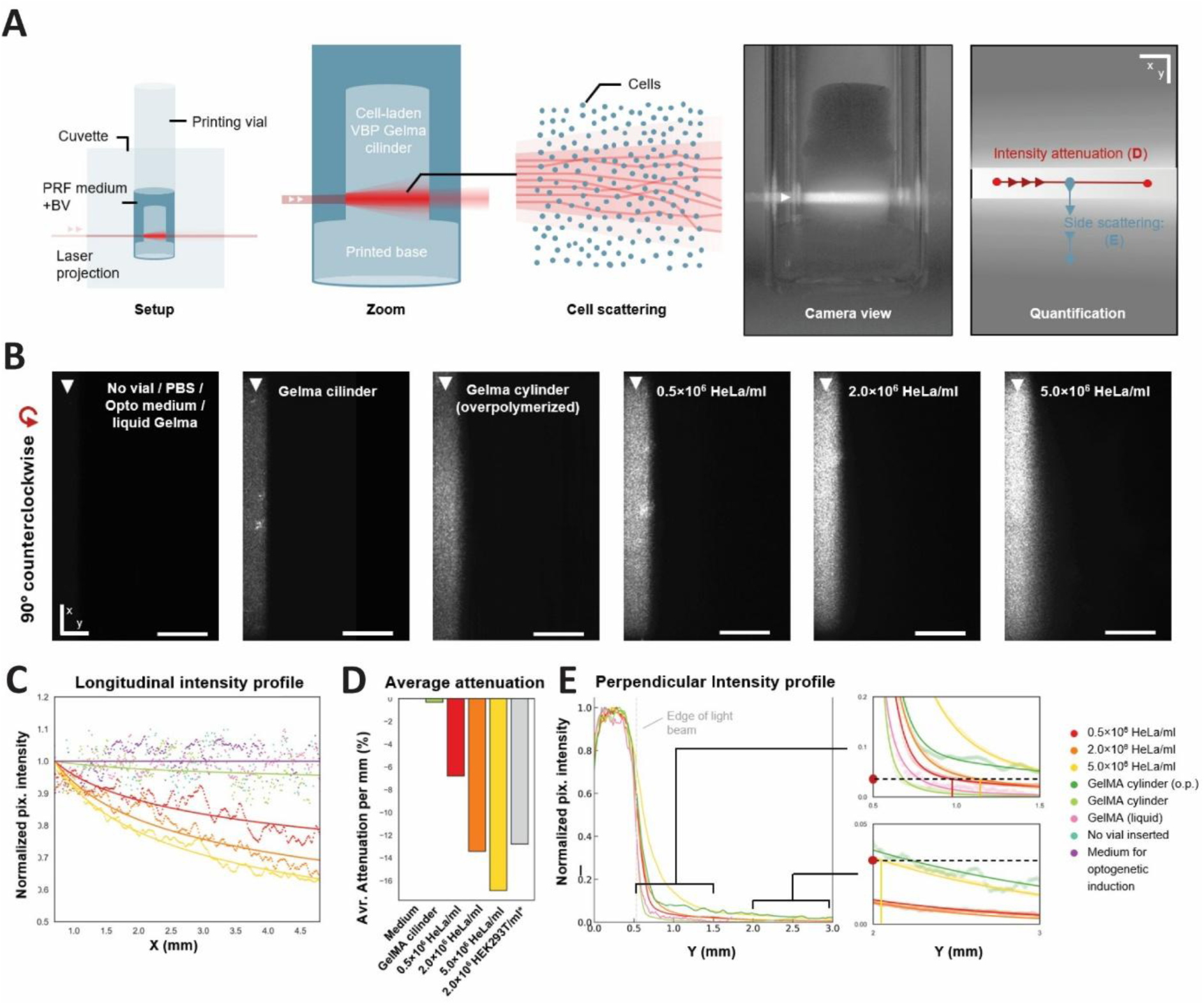
Scattering profiles of direct NIR light projections in volumetrically printed cell-laden GelMA constructs. (A) Schematic of the projection setup used for measuring scattering profiles. i: The printing vial was submerged in demineralized water inside a cuvette. The volumetrically printed construct was positioned in phenol red– free medium supplemented with 25 μM BV. A 0.532 mm × 4 mm laser beam was projected through the cuvette onto the target region within the printing vial. ii: Zoom view of an example cell-laden cylindrical construct, standing on a base, submerged with the medium, including a hypothetical scattered light profile. iii: Schematic of Mie scattering of the light by the cells in the print. (B) i: Contrast-reduced preview from the IR-camera side perspective showing projections passing through the construct. (ii) Schematic of the measured profiles: longitudinal intensity (shown in panel D) and perpendicular light diffusion (shown in panel E). (C) Raw images of projections passing through different constructs, rotated 90° counterclockwise. S.B. = 1 mm (D) i: Beam attenuation measured by pixel intensity normalized to start at 0 at x = 0.7 mm to eliminate edge artefacts. Exponential decay curves were fitted to condition average profiles (n = 2). ii: Average percent light attenuation per mm travelled through the construct, derived from decay fit curves. (E) Perpendicular intensity profiles measured by normalized pixel intensity. Grey dotted line at y = 0.532 represents the edge of the projection. ii and iii are zoom-ins of i. ‘▲’ indicates laser input location. Avr. = Average, o.p. = overpolymerized, Pix. = pixel, PRF = phenol-red free.

**Supplementary Figure S16.**
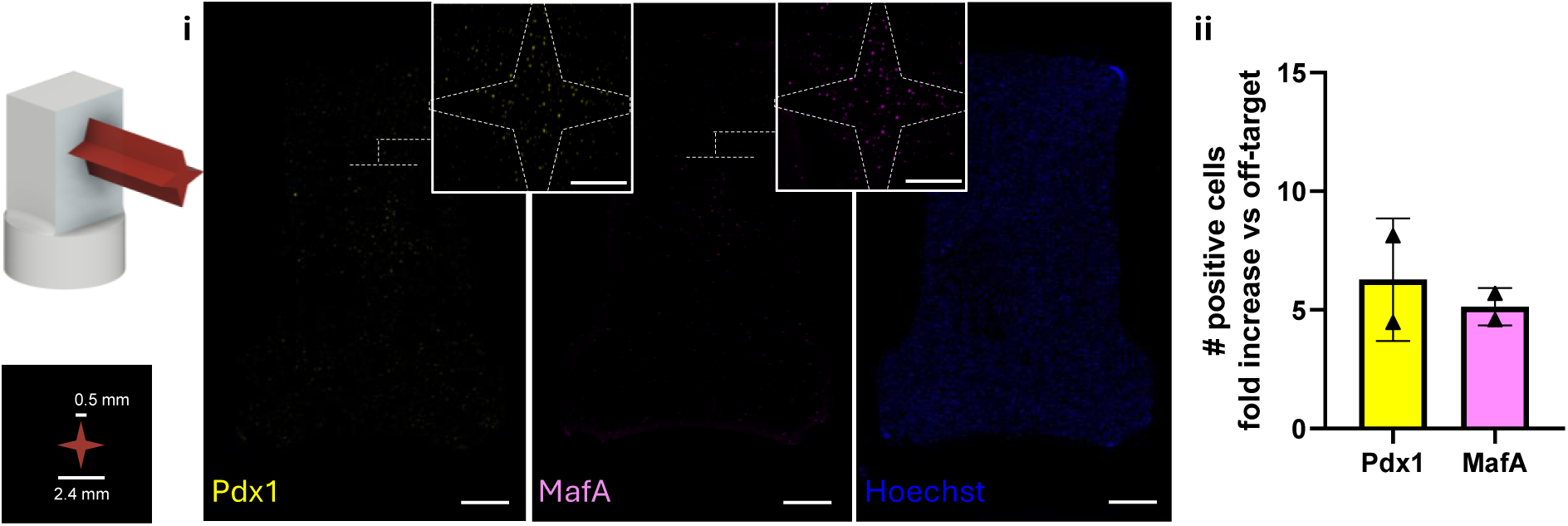
Pilot experiments of spatially patterned optogenetic stimulation using NIR-light projections from the bioprinter. Projection of a star pattern (arm width 0.5 mm, axis length 2.4 mm) into bioprinted constructs enables patterning of gene expression for (i) Pdx1 and MafA with low signal in off-target areas, and higher signal on-target. Although off-target activation of cells outside the projected pattern was also observed, particularly close to the border of the hydrogels, were scattering is accentuated by the transition from the culture media to the GelMA, (ii) zonally confined optogenetic stimulation successfully resulted in a 6.28 ± 2.58 (Pdx1) and 5.12 ± 0.79 (MafA) fold increase in the density of activated cells within the star pattern. The overall density of higher signal cells in the on-target areas is approximately 5-fold compared to the off-target region (N = 2). Scale bar in F is 1 mm, inset is 500 µm.

